# Inference of selection from genetic time series using various parametric approximations to the Wright-Fisher model

**DOI:** 10.1101/696955

**Authors:** Cyriel Paris, Bertrand Servin, Simon Boitard

**Affiliations:** Génétique, Physiologie Systèmes d’Elevage, INRA, Centre Occitanie-Toulouse, 31326 Castanet-Tolosan, France

## Abstract

Detecting genomic regions under selection is an important objective of population genetics. Typical analyses for this goal are based on exploiting genetic diversity patterns in present time data but rapid advances in DNA sequencing have increased the availability of time series genomic data. A common approach to analyze such data is to model the temporal evolution of an allele frequency as a Markov chain. Based on this principle, several methods have been proposed to infer selection intensity. One of their differences lies in how they model the transition probabilities of the Markoiv chain. Using the Wright-Fisher model is a natural choice but its computational cost is prohibitive for large population sizes so approximations to this model based on parametric distributions have been proposed. Here, we compared the performance of some of these approximations with respect to their power to detect selection and estimation of the selection coefficient. We developped a new generic Hidden Markov Model likelihood calculator and applied it on genetic time series simulated under various evolutionary scenarios. The Beta-with-Spikes approximation, which combines discrete fixation probabilities with a continuous Beta distribution, was found to perform consistently better than the others. This distribution provides an almost perfect fit to the Wright-Fisher model in terms of selection inference, for a computational cost that does not increase with population size. We further evaluate this model for population sizes not accessible to the Wright-Fisher model and illustrate its performance on a dataset of two divergently selected chicken populations.

## Introduction

Detecting the molecular basis of adaptation in natural species is one of the major questions in population genetics. Most studies in this context rely on contemporary genomic data and exploit several typical signatures left by the spread of a benefical allele in a population: decreased heterozygosity, extended linkage disequilibrium (LD), increased differentiation with other populations …, see Vitti et al. (2013) for a review.

Genomic time series provide an even more direct access to past allele frequency changes, allowing a more precise estimation on the onset or the intensity of selection. With the combined advances in sequencing technologies, ancient DNA processing and cryoconservation, this type of data are becoming accessible to many researchers and are found in various contexts. On a long time scale, time series of ancient DNA have been used for example to detect the genetic basis of adaptation in humans from Eurasia (Mathieson et al., 2015) or to infer allele frequency trajectories at several causal variants or quantitative trait loci in domestic horses (Fages Antoine et al., 2019). On a shorter time scale, genetic time series are provided by the sequencing of samples conserved in biobanks during decades. This approach allowed for instance to assess the recent evolution of genetic diversity in Holstein cattle and to predcit the selective value that could be obtained by using semen from ’historical’ bulls in this breed (Doekes et al., 2018). Controlled evolution experiments also provide a lot of interesting temporal genetic data and concern a large set of organisms. Such experiments generally imply intentional selection for a given trait of interest, e.g. thermal tolereance in drosophila (Tobler et al., 2015) or intra-muscular ultimate pH in chicken (Bihan-Duval et al., 2018).

Following the increased production of genetic time series, several methods allowing to exploit such data for selection inference have been developed in the two last decades (Malaspinas, 2015). Most of them infer selection at a given variant based on temporal observations at this variant only, i.e. without exploiting LD information (but see (Terhorst et al., 2015; He et al., 2019)). In this context, several statistical approaches have been proposed to model the link between the full temporal trajectory of a population allele frequency, which is informative about selection intensity at the locus, and the (partial) genetic samples observed at some specific dates along this trajectory. These approaches include approximate bayesian computation (ABC) (Foll et al., 2015; Sackman et al., 2019), bayesian path augmentation Schraiber et al. (2016) or hidden Markov models (HMM) (Bollback et al., 2008). The HMM approach arises quite naturally when analyzing genetic time series, as it accounts both for the Markovian property of allele frequency trajectories over time and for the fact that these trajectories are latent variables. Besides, efficient algorithms have been developped for likelihood evaluation and parameter estimation in such models (Rabiner, 1989). Following this approach, one important choice concerns the modelling of the underlying allele frequency trajectory at a locus. In the initial work of Bollback et al. (2008), this process was modelled by the Wright-Fisher (WF) diffusion and the partial differential equation (PDE) resulting from this diffusion between each pair of consecutive sampling times was solved using a numerical scheme. More recently, Song and Steinrücken (2012) derived an analytical solution of this PDE based on an infinite series expansion and integrated this solution into the HMM framework to detect selection on ancient DNA (Steinrücken et al., 2014). The difusion process can also be approximated using Markovian continuous time processes with a relatively low number of discrete spaces (Malaspinas et al., 2012; Ferrer-Admetlla et al., 2016). All these approaches are quite computationally demanding, moreover the WF process converges to the diffusion only for weak selection (Ewens, 2004).

Considering the WF model as a reference, an even more natural approach is to compute transition probabilities of the Markov chain using this model directly (Iranmehr et al., 2017; Hubert et al., 2018). However, this implies that the dimension of the transition matrix is equal to the population size so this approach is in practice only possible for small populations. A more efficient strategy is to approximate the WF process by a parametric continuous distribution, whose first two moments fit those of the WF and whose transitions can be computed using a simple analytical formula. This moment-based approximation was implemented by Lacerda and Seoighe (2014) and Terhorst et al. (2015) using a Gaussian distribution and by Gompert (2016) using a Beta distribution. However, Tataru et al. (2016) showed that these two distributions accurately approximate the WF only for very short time scale and weak selection and that a much closer fit can be obtained by incorporating fixation probabilities in the Beta distribtuion, leading to what they called the Beta with Spikes distribution. They illustrated the performance of this distribution to infer branch lengths in a population tree, but did not consider the potential application for selection inference.

Here we compare the performance of these different approximations for inferring selection. For this purpose, we implemented a generic HMM allowing to estimate selection intensity by maximum likelihood and to detect selected loci by a likelihood ratio test. Transition probabilities in this model can be computed using either the WF model, the Gaussian approximation, the Beta approximation or the Beta with Spikes approximation. We evaluate the quality of inference provided by these different transitions in a large range of selection scenarios and show an application of the HMM approach to a dataset concerning the experimental divergent selection of two chicken lines (Bihan-Duval et al., 2018).

## Methods

### Statistical models for genetic time-series

#### Hidden Markov Model framework

Let *A* be a locus of interest accepting two alleles: *A*_0_ is called the (arbitrary) *reference allele* while *A*_1_ is called the *alternative allele*. Considering a diploid organism, we assume each genotype is associated to a particular fitness: the homozygote reference allele fitness is 1, the heterozygote form fitness is 1 + *sh* and the homozygote alternative allele fitness is 1 + *s*, where *s* is the selection parameter and *h* the dominance parameter. Assume a population has been sampled at *n* different dates *t*_1_ = 0, …, *t*_*n*_ = *T* (in generations after the first sampling date) in order to assess the evolution of the frequency of allele *A*_1_. We note *X*_(*t*)_ the true frequency of *A*_1_ at date *t*. At date *t*_*k*_ (*k* ∈ {1, …, *n*}), we note 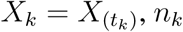, the total number of alleles sampled and *Y*_*k*_ the number of *A*_1_ alleles sampled. Finally, we note *N*_*e*_ the haploid effective size of the population.

In this study we will follow a common approach to model such data, first proposed by Bollback et al. (2008) and use a Hidden Markov Model (HMM, see Cappé et al. (2005) for a general de-scription). This HMM (Figure 1) is a bivariate Markov process (*X*_*k*_, *Y*_*k*_)_*k*≥1_ with *X*_*k*_ ∈ [0, 1] and *Y*_*k*_ ∈ {0, …, *n*_*k*_}. (*X*_*k*_) is a Markov process with *transition kernel* **Q**_***k***_ between *X*_*k*−1_ and *X*_*k*_. Given (*X*_*k*_), observations are independant and the *emission kernel* of *Y*_*k*_ given *X*_*k*_ is noted *g*_*k*_. The distribution of *X*_1_ is noted *ν*. This model is fully specified by setting *ν, Q*_*k*_ and *g*_*k*_, and each of them can be thought of as a probability:

**Figure 1:**
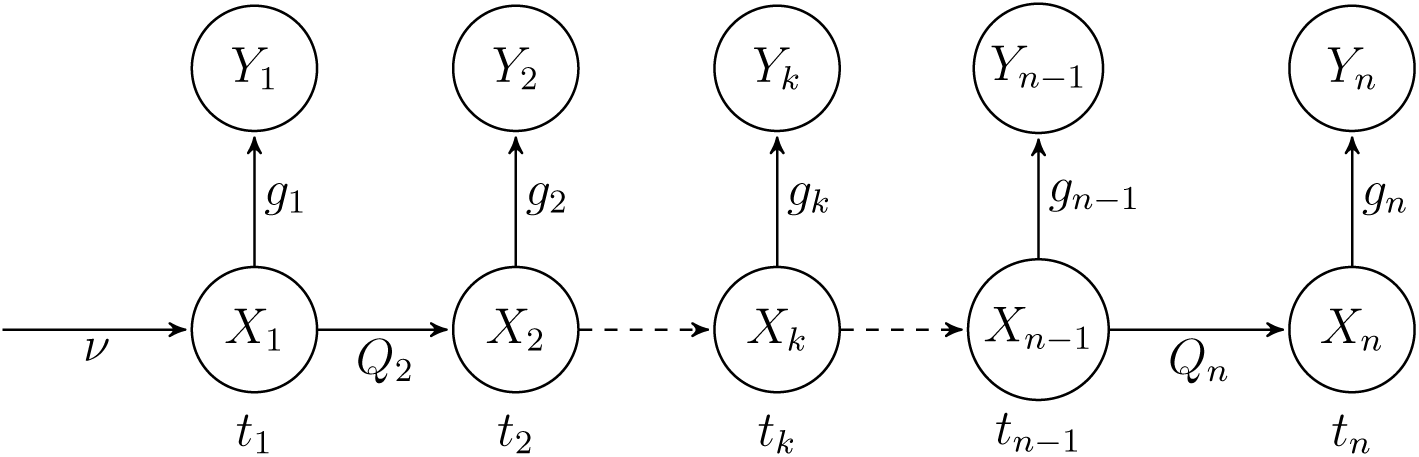
Graphical representation of the Hidden Markov Model for genetic time series.

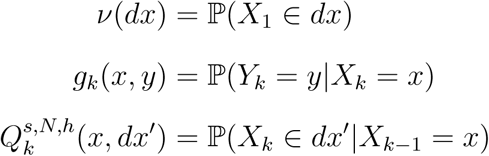

In this study we assume that the distribution of *X*_1_ is uniform on [0, 1] (*ν*(*dx*) = *dx*). As the observations *Y*_*k*_ are sampled from a population with allele frequency *X*_*k*_, the distribution of *Y*_*k*_ given *X*_*k*_ is a binomial distribution with parameters *n*_*k*_ and *X*_*k*_:

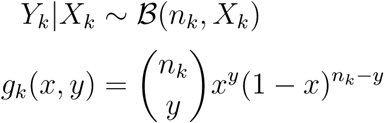

For a given *x, Q*_*k*_(*x*, .) is the distribution of *X*_*k*_ given that the previous frequency *X*_*k*−1_ is *x*. Note that the Markov chain *X*_*k*_ is typically not homogeneous as for example the duration between successive sampling dates may vary. Apart from the time between samples, *Q*_*k*_ also depends on the transition model and on parameters *s, N*_*e*_ and *h*. One of the main objective of this study is to compare different possible transition models which are described below.

In this HMM framework, the likelihood is:

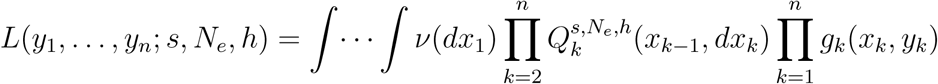

A direct calculation of this likelihood is very costly because the integral is over [0, 1]^*n*^. The usual way to tackle this problem is to introduce the forward measure *α*_*k*_, which is the joint probability of the *k* first observations and the *k - th* state:

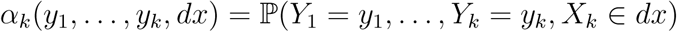

It is computed using the following recursion:

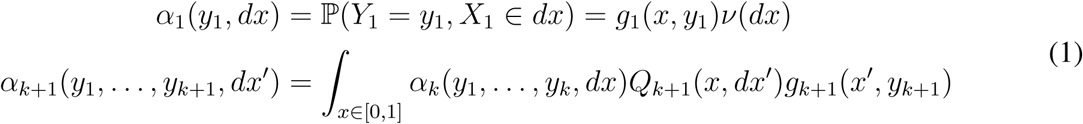

The likelihood is obtained by integrating the last forward measure *α*_*n*_ over all possible states *X*_*n*_, using the following relation:

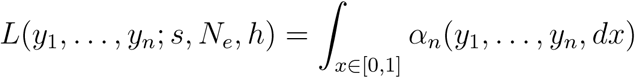

Using this approach, the likelihood calculation involves only *n* integrals over [0, 1] instead of one integral over [0, 1]^*n*^. For a later use, let’s also define the log likelihood:

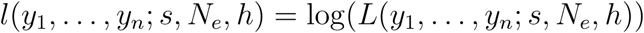

This likelihood forms the basis of statistical inference on the population parameters *N*_*e*_, *s* and *h*. In this study, we focus on the inference on the selection parameter *s* and will therefore assume *N*_*e*_ and *h* to be known. We estimate the selection parameter *s* by maximum likelihood:

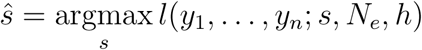

To build a test for the null hypothesis (*s* = 0), we use the log likelihood ratio statistic:

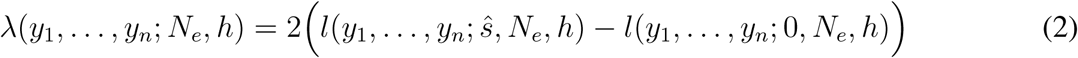

Under the null, *λ* asymptotically follows a *χ*^2^ with one degree of freedom, a property that allows to derive p-values which are useful quantities in particular in a multiple testing context.

Having described the general framework used in this study, we now turn to its implementation that requires specifying the transition kernel *Q*_*k*_(*x*, .). One objective of this work was to investigate different models for this transition kernel. A reference model in this context is the one-locus Wright-Fisher (WF) model under diploid selection pressure (Ewens, 2004). However, this model has computational limitations and, in the HMM famework, must generally be approximated. We first present the WF model and then several continuous approximations of this model based on a common approach: the method-of-moments.

#### Wright-Fisher transition model

The WF model is a discrete time model that assumes random mating and non overlapping generations. Let *N*_*e*_ be the haploid (constant) effective population size. Under this model, *X*_(*t*)_ can take a discrete number of values in 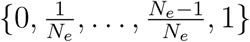 and the transition kernel of this process is a transition matrix with size (*N*_*e*_ + 1) × (*N*_*e*_ + 1). Let **P** designate the transition matrix of this process in one generation. Then, 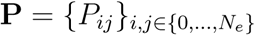 and

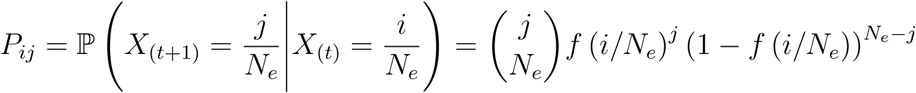

where *f* is the diploid fitness function (Ewens, 2004):

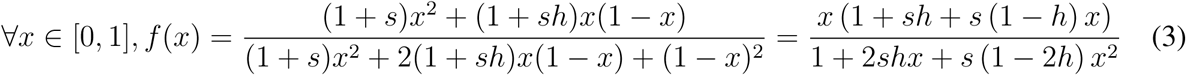

Note that this function can easily incorporate the possibility of mutations (Ewens, 2004) but it is not considered here for the sake of simplicity.

Given this one step transition matrix, the transition matrix of the HMM is obtained by taking **P** to the required power 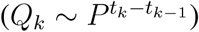:

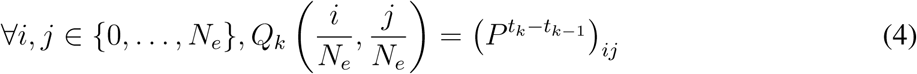

In the WF model, the effective population size *N*_*e*_ is a critical parameter affecting computational properties. More precisely, the transition matrix memory occupation is 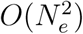 and the forward algorithm computation time is at least 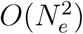. Moreover, the transition matrix **Q**_***k***_ is obtained by taking a (*N*_*e*_ + 1) × (*N*_*e*_ + 1) matrix to the power *t*_*k*_−*t*_*k*−1_, with a complexity in 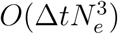. It can thus be very costly for large *N*_*e*_ and large intersample time. To overcome this issue, one approach is to approximate the Wright-Fisher process with a continuous space process such that integral calculations of the forward algorithm are made over a discretization of [0, 1], with a grid size (i.e. a number of discretization points) much smaller than *N*_*e*_. We followed this approach here and approximated the WF process using the method-of-moments.

#### Parametric approximations based on the method of moments

The principle of the method-of-moments consists in modelling the transition kernel *Q*_*k*_ with a parameteric distribution, matching its mean and variance to those of the WF model. The advantages of using parameteric distributions are (i) to be able to compute transitions faster than rising the transition matrix at the good power, (ii) use smaller transition matrix in the forward algorithm. This approach was first proposed by Lacerda and Seoighe (2014) with a Gaussian distribution and was then used in several other studies (Terhorst et al., 2015; Tataru et al., 2016; Gompert, 2016). One prerequisite of the method-of-moments is to know the moments of a WF model for any value of *s, h* and *N*_*e*_. Under selection (*s*≠0) no analytical formula of these moments exists but they can be approximated using a recursion inspired by (Lacerda and Seoighe, 2014; Terhorst et al., 2015; Tataru et al., 2016). In the WF model, moments at generation *t* + 1 can be derived from those at generation *t*:

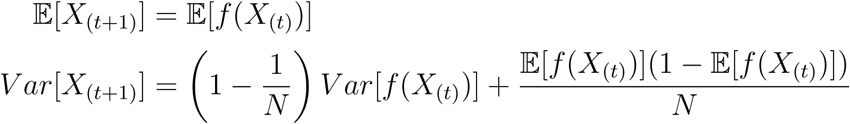

For models with selection, the fitness function *f* (3) is non affine so this recursion cannot be solved analytically. However, using a Taylor expansion of *f* around 𝔼[*X*_(*t*)_], it is possible to build a recursion allowing to approximate 𝔼[*X*_(*t*)_] and *V ar*[*X*_(*t*)_]. Let *µ*_*t*_ and 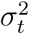 be these approximations at generation *t*, they can be computed as follows (see Supplemental File S1 for details):

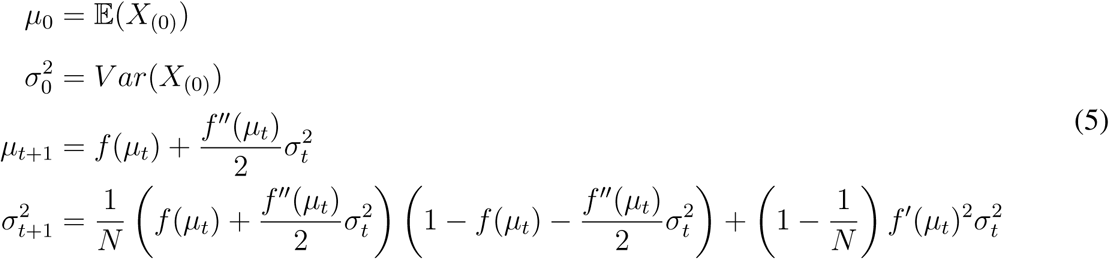

Note that as f is a rationale fraction, its derivatives can be calculated exactly. To simplify notations in the following, we further define 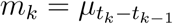 and 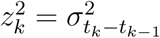.

Having established approximations to the Wright-Fisher moments in the general case, different parametric distributions can be chosen to approximate the transition kernel. We shown Figure **??** examples of approximations using models presented below.

#### Gaussian Model (Ga)

The first possibility, considered in the original description of the method-of-moments approach (Lacerda and Seoighe, 2014) is to use the Gaussian distribution (Nicholson et al., 2002). Indeed, when 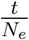 and *N*_*e*_*s* are small, the rescaled Wright-Fisher process converges to a Gaussian process and the HMM transition kernel is:

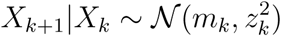

One of the main issues with the gaussian model is that *X*_(*t*)_ can take values outside [0, 1]. To tackle this issue we chose to reject values distributed outside [0, 1] and appropriately rescale the Gaussian density. Another alternative (not considered here) is to allocate the mass of the distribution outside [0, 1] to discrete fixation probabilities at 0 (*i.e. P* (*X*_(*t*)_ = 0) := *P* (*X*_(*t*)_ ≤ 0)) and 1 (*i.e. P* (*X*_(*t*)_ =1) := *P* (*X*_(*t*)_ ≥ 1)) (Nicholson et al., 2002).

#### Beta Model (Be)

A second possible model is the Beta model (Tataru et al., 2016; Hui and Burt, 2015; Gompert, 2016). Use of this model can be motivated by the fact that it is adequately distributed on [0, 1] and because the stationary distribution of the diffusion approximation under neutrality is a Beta distribution. In addition, the family of Beta distributions encompasses a lot of different shapes that can fit many behaviors for allele frequency variation. In this model, the HMM transition kernel can be written as:

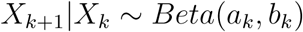

with:

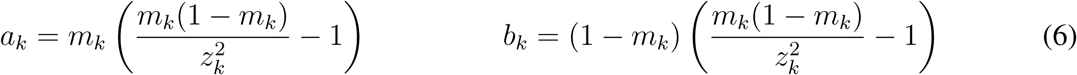

#### Beta with spikes Model (BwS)

We finally considered the beta with spikes model described by Tataru et al. (2016). In contrast to the two continuous models described above, this model explicitely accounts for the possibility of allele fixation: it is a mixture of a beta distribution and two discrete fixation probabilities at 0 and 1. According to Tataru et al. (2016), this model is a better approximation of the Wright-Fisher than the Ga and Be models.

Let *p*_0,*k*_(*x*) = ℙ(*X*_*k*_ = 0|*X*_*k*−1_ = *x*) and *p*_1,*k*_(*x*) = ℙ(*X*_*k*_ = 1|*X*_*k*−1_ = *x*). The transition kernel is:

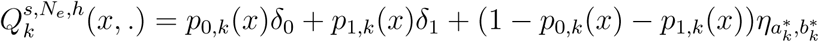

where 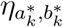 is a beta distribution describing the transition from *X*_*k*_ to *X*_*k*+1_ conditional on the fact that *X*_*k*+1_ did not fix in 0 or 1. Parameters 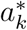 and 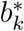 are obtained with a formula similar to (6), but based on the conditional (rather than absolute) moments of the WF process:

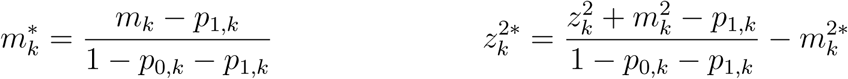

In the beta with spikes model, fixation probabilities must be computed in addition to WF moments. Let us define *p*_0,*t*_ = ℙ(*X*_(*t*)_ = 0) and *p*_1,*t*_ = ℙ(*X*_(*t*)_ = 1). We used the following recursion, proposed by Tataru et al. (2015) assuming beta with spikes transitions:

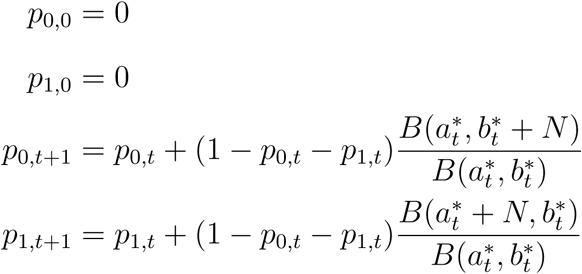

where *B*() denotes the Beta function.

#### Implementation

In this study, the likelihood was always evaluated on a grid of *N*_*e*_*s* composed of 600 values ranging from *-*100 to 300. For WF transitions the integrals involved in the likelihood computation using the forward algorithm (1) are finite sums. However for the parametric approximations to the WF these integrals are continuous and must therefore be numerically approximated. Here, this was done using the trapezoidal rule on an integration grid composed of 129 points evenly spaced on [0, 1]. Note that the calculation of these transitions is only required once per dataset. This implementation of the HMM is freely available at https://github.com/CyrielParis/compareHMM.

### Comparison of HMM transition models

We studied the quality of the transition approximation to the WF process. For this objective, we set the effective population size to *N*_*e*_ = 100 haploids to render the comparisons computationally tractable. We compared the Ga, Be and BwS models as approximations to the Wright-Fisher process, first by evaluating their fit to the WF distribution and second by measuring their impact on selection inference.

#### Approximation of the WF transition

We first studied the quality of approximations of the WF transition kernel. For all parametric distributions, at least part of the WF discrete transition is approximated by a continuous density. Thus, measuring the fit of these approximations requires comparing continuous densities to discrete probabilities. In such a case, classical distances aimed at comparing densities, such as the Kullback-Liebler divergence or the Hellinger distance are not valid. Therefore we used the *Wasserstein distance* defined as the *L*_1_ distance between the cumulative distribution functions of two distributions and which is well defined in all cases considered here. The fit of the Ga, Be and BwS models were evaluated over the parameter set *h* = 0.5, (*t*_1_ −*t*_0_) ∈ [1, 20], *N*_*e*_*s* ∈ {0, 10, 100}, and starting allele frequency covering all *i/N*_*e*_ values for *i* ∈ {0, …, *N*_*e*_}

#### Consequences for statistical inference

We computed the likelihood profile (see implementation paragraph) for genomic time series randomly simulated with the Wright-Fisher process. We set the initial allele frequency to 0.1 or 0.5 and its selection parameter varying from 0 (neutral case) to 1 (strong selection). We simulated a random sampling of *n*_*k*_ = 30 alleles at 10 evenly spaced dates over a total range of *T* ∈ {9, 45, 90, 180} generations (*i.e. t*_*k*_ −*t*_*k*−1_ ∈ {1, 5, 10, 20}). We ran 100000 simulations for *s* = 0 and 10000 for other *s* (see Figure S2). To compare the Ga, Be and BwS models in terms of selection inference, we focused on four summarizing quantities computed from the likelihood: the log-likelihood value at *s* = 0, the maximum likelihood estimator ŝ, the log-likelihood value at *s* = ŝ, and the log-likelihood ratio statistic (2).

### Inferring selection using the HMM

As the BwS model was found to be a good approximation to the Wright-Fisher (see Results), we further studied its statistical properties on simulations for higher *N*_*e*_ and applied it to detect selection on a real data set.

#### Simulation study

To study the statistical properties of the BwS model, we created new datasets designed to represent consistent selection dynamics as for the *N*_*e*_ = 100 dataset but with larger *N*_*e*_. To do so, we considered that the diffusion approximation to the WF suggests that selection dynamics are driven by the two compound parameters *N*_*e*_*s* and *T/N*_*e*_. Therefore the new datasets were created by increasing *N*_*e*_ to 1,000 and 10,000 but in each case adjusted *T* and *s* so as to keep the same set of *N*_*e*_*s* and *T/N*_*e*_ values as for the *N*_*e*_ = 100 dataset. Specifically we considered *N*_*e*_*s* ∈ [0, 100] and *T/N*_*e*_ ∈ {0.09, 0.45, 0.9, 1.8}. We also kept the same sampling pattern of *n*_*k*_ = 30 haploids sampled at 10 evenly spaced dates over the total range *T* and an initial allele frequency of 0.1 or 0.5.

#### Real data analysis

We applied our method using the BwS model on a dataset of two experimental populations of chicken divergently selected for the intra muscular ultimate pH (pHu) (Bihan-Duval et al., 2018; Alnahhas et al., 2016). This selection experiment was conducted for 5 generations and at each generation (including the starting population) samples were collected and genotyped for 40,199 markers. Sample sizes in each line (pHu- and pHu+) are given in Table 1. Observed allele frequencies in the two lines, for all markers and generations, are provided in File S2 at plink format. For our analysis we considered the two lines as two different populations.

**Table 1:** Haploid sample sizes across 6 generations of chicken populations selected for intra-muscular ultimate pH.

| Dates (in generations) | 0 | 1 | 2 | 3 | 4 | 5 |
| --- | --- | --- | --- | --- | --- | --- |
| pHu+ | 102 | 28 | 42 | 56 | 54 | 60 |
| pHu- | 102 | 34 | 46 | 44 | 54 | 56 |

Our current implementation of the method does not estimate the *N*_*e*_ parameter which is not known in this real dataset. To remedy this issue, we estimated *N*_*e*_ in each line using the NB R package (Hui and Burt, 2015). Motivation for using this method to estimate *N*_*e*_ comes from the fact that the underlying model is a HMM with beta transitions, similar to our general framework. To estimate *N*_*e*_, we removed trajectories starting near edges (frequencies below 20% or above 80%), running the analysis over 27,669 loci.

We removed from the analysis all trajectories fixing in one step as we used the *χ*^2^(1) distribution to compute p-values. SNP were called significant with a FDR threshold of 5% estimated using the q-value R package (Storey et al., 2015).

## Results

We have established a general HMM framework that allows to perform statistical inference from genetic time-series data. Within this framework, different transition models can be used and easily compared. In this section we will first present results comparing the quality of approximations of the Gaussian (Ga), Beta (Be) and Beta-with-spikes (BwS) transition kernels on their ability to approximate the Wright-Fisher (WF) kernel. Next, we study in more details the statistical performances of the BwS for detecting selection and estimating its intensity. Finally, we illustrate how the BwS model can be used on real data by re-analyzing an experimental evolution dataset.

### Approximation of the Wright-Fisher model

#### Approximation of the WF transition

We first compared each continuous approximation with the WF using the Wasserstein distance. Figure 2 shows for each model and for different selection intensities the heatmap of the Wasserstein distance between the continuous approximation and the true WF distribution. In each panel, the starting allele frequency is on the x axis and the number of generations between two dates is on the y axis. Each cell is colored according to the Wasserstein distance. In principle this distance ranges from 0 which is a perfect match to 1 which is, for an allele frequency, the worst possible distance. For this set of parameters, the Wasserstein distance ranged from 0 to ≈ 0.5. According to this distance, all approximations are valid for intermediate starting allele frequencies and a small number of generations but the accuracy of all approximations decreases as the number of generations increases. The accuracy of the approximation also worsens when the selection parameter increases, especially for small starting allele frequency. Generally these results show that the Be and BwS models are better approximations than the Ga model1, the BwS model being overall slightly better than others. These results are consistent with those obtained by Tataru et al. (2016) although they used a different approximation for the moments and a different measure of the distance between distributions.

**Figure 2:**
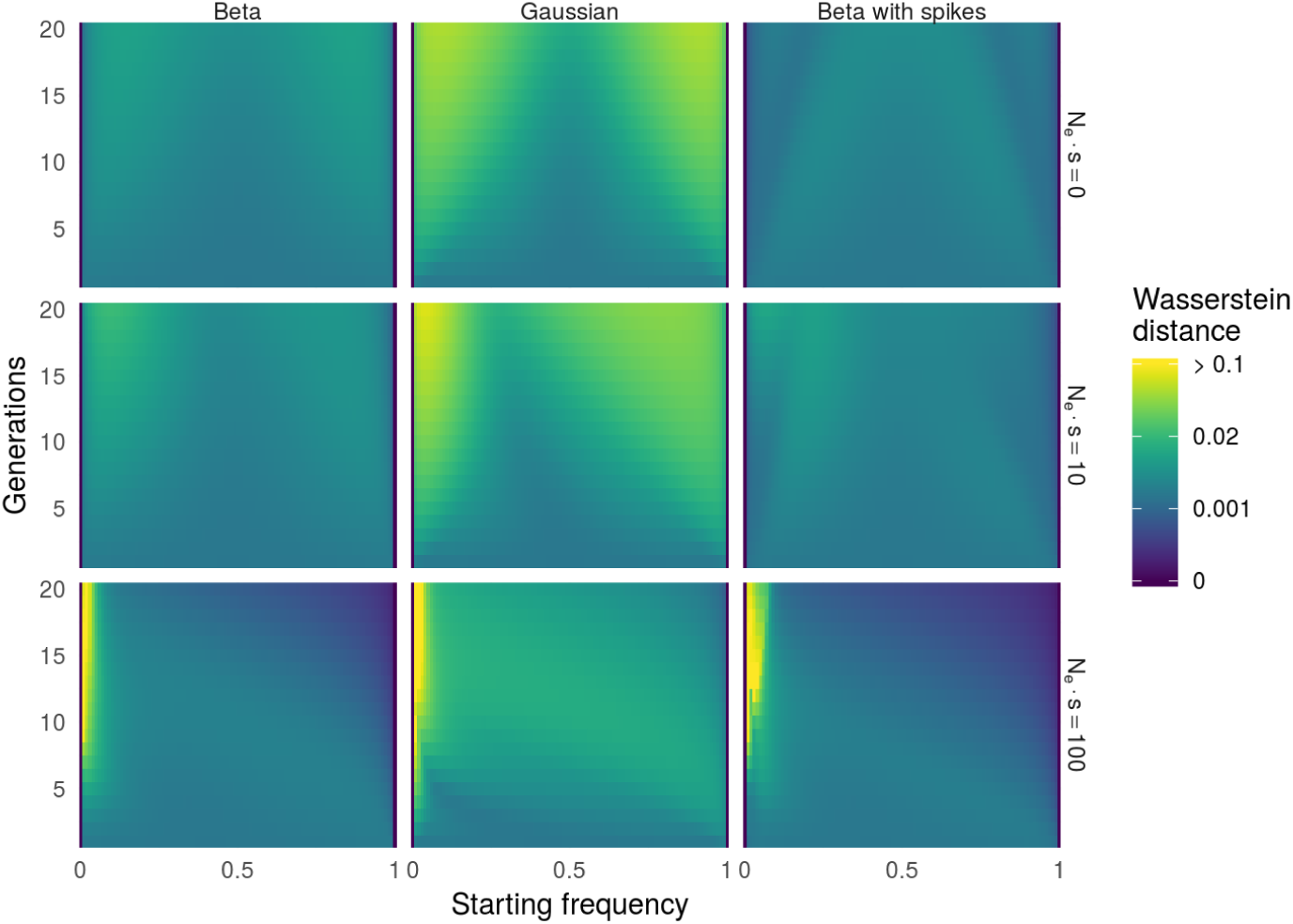
Wasserstein distances to Wright-Fisher transition for three continuous approximations and for varying starting allele frequencies and durations. Top panel: Neutral evolution (*N*_*e*_*s* = 0). Middle panel: Mild selection (*N*_*e*_*s* = 10). Bottom panel: Strong selection (*N*_*e*_*s* = 100)

The error in the approximation can come from the model itself but also from the use of approximated moments. To evaluate what proportion of the distance to the true distribution comes from the moment approximation, we computed the Wasserstein distance of each continuous distribution using the true WF moments (Equation 4, Figure S3). Figure 3 presents this proportion for the BwS model under neutrality (left) and under selection (middle *N*_*e*_*s* = 10 and right, *N*_*e*_*s* = 100). Under neutrality the error comes mainly from the continuous approximation. This is also usually the case under selection although for small starting frequencies (*x*_1_ *<* 0.1) and large duration between samples the error in the approximation of moments is the most influential.

**Figure 3:**
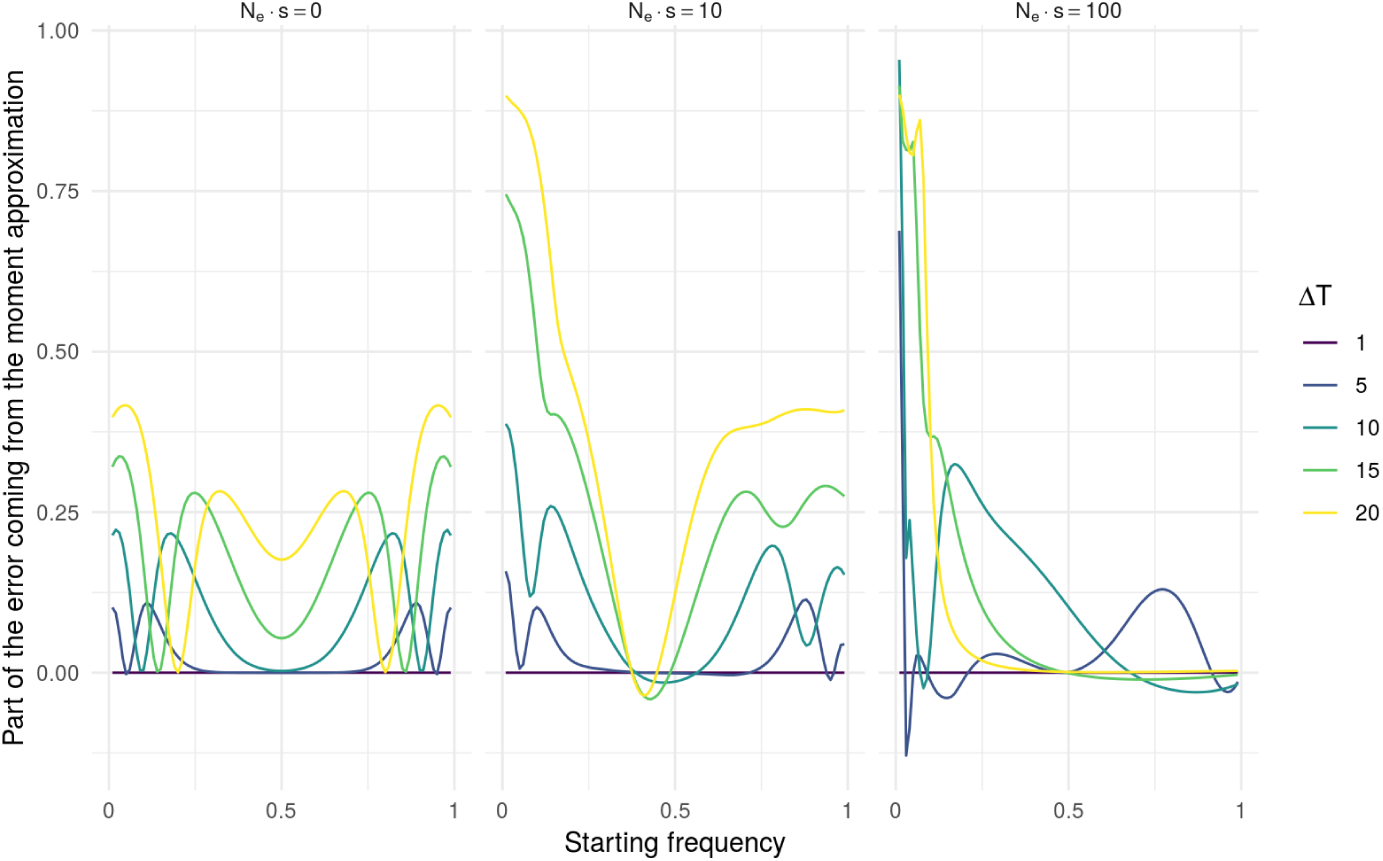
Proportion of the Wasserstein distance of the BwS due to the approximation of moments. Negative values correspond to very rare cases where the transition with approximated moments is better than with true moments.

#### Approximation of the WF likelihood

Next, we evaluated the ability of continuous WF approximations to recover the WF likelihood of genetic time series data. To do so, we simulated trajectories under different evolutionary scenarios (see Methods) and compared the inference obtained with each continuous model to the one obtained using the Wright-Fisher model.

Simulations of time-series genetic data to evaluate the quality of parametric approximations to the Wright-Fisher were performed under this reference model. Hence, using the HMM with WF transitions should provide the best statistical behavior and be considered a reference inference method. Before assessing the quality of approximations to the WF model with respect to their accuracy for computing the WF-based likelihood, it is necessary to ensure that this model itself provides sensible likelihood inference. Results on simulated data showed that this was not always the case: a family of trajectories were found to lead to unreliable results even when using the WF model for inference. These trajectories correspond to cases where the observed allele frequency reached fixation in one time step (*i.e.* ∀*k* > 1, *Y*_*k*_ = 0 or ∀*k* > 1, *Y*_*k*_ = *n*_*k*_). In such cases the likelihood function is monotonic in *s*: the *s* value that maximizes the probability of fixation at 1 in one time step is *s* = ∞ and similarly the *s* value that maximizes the probability of fixation at *s* = 0 in one time step is *s* = *-*1, essentially corresponding to an infinite *s* for the other allele. In these cases, likelihood inference is not valid. They could be deemed merely pathological but they occur with a non negligible probability that is increasing as the intersample duration or the selection intensity increases. In the following, we excluded such cases when evaluating the statistical properties of inference models. However we studied their occurrence frequency (see below).

Figure 4 presents four different statistics based on the likelihood of simulated data under various scenarios (see Methods), comparing results obtained for each parametric approximation to those obtained using the WF: Figure 4a presents results on the likelihood itself at *s* = 0, Figure 4b the maximum likelihood estimates (MLE), Figure 4c the likelihood at the MLE (which can be different from the WF one) and Figure 4d the likelihood ratio statistic. For all these statistics and independently of the scenario simulated, the BwS and the WF provide essentially the same inference. Only for the MLE a few cases exist where the two transition model disagree (Figure 4b). However, this is never true for the LMLE (Figure 4c) so when the two models disagree for ŝ they still give similar likelihood values at the MLE. Overall this shows that the few large differences in MLE between BwS and WF most likely correspond to cases where the data are not very informative on the selection intensity. On the contrary, there are many scenarios where the Be and Ga models have very different values from the WF ones for all statistics considered. This shows that in many scenarios they are bad approximations to the WF model. So the BwS model is strictly better than the Ga and Be in terms of approximating the Wright-Fisher likelihood.

**Figure 4:**
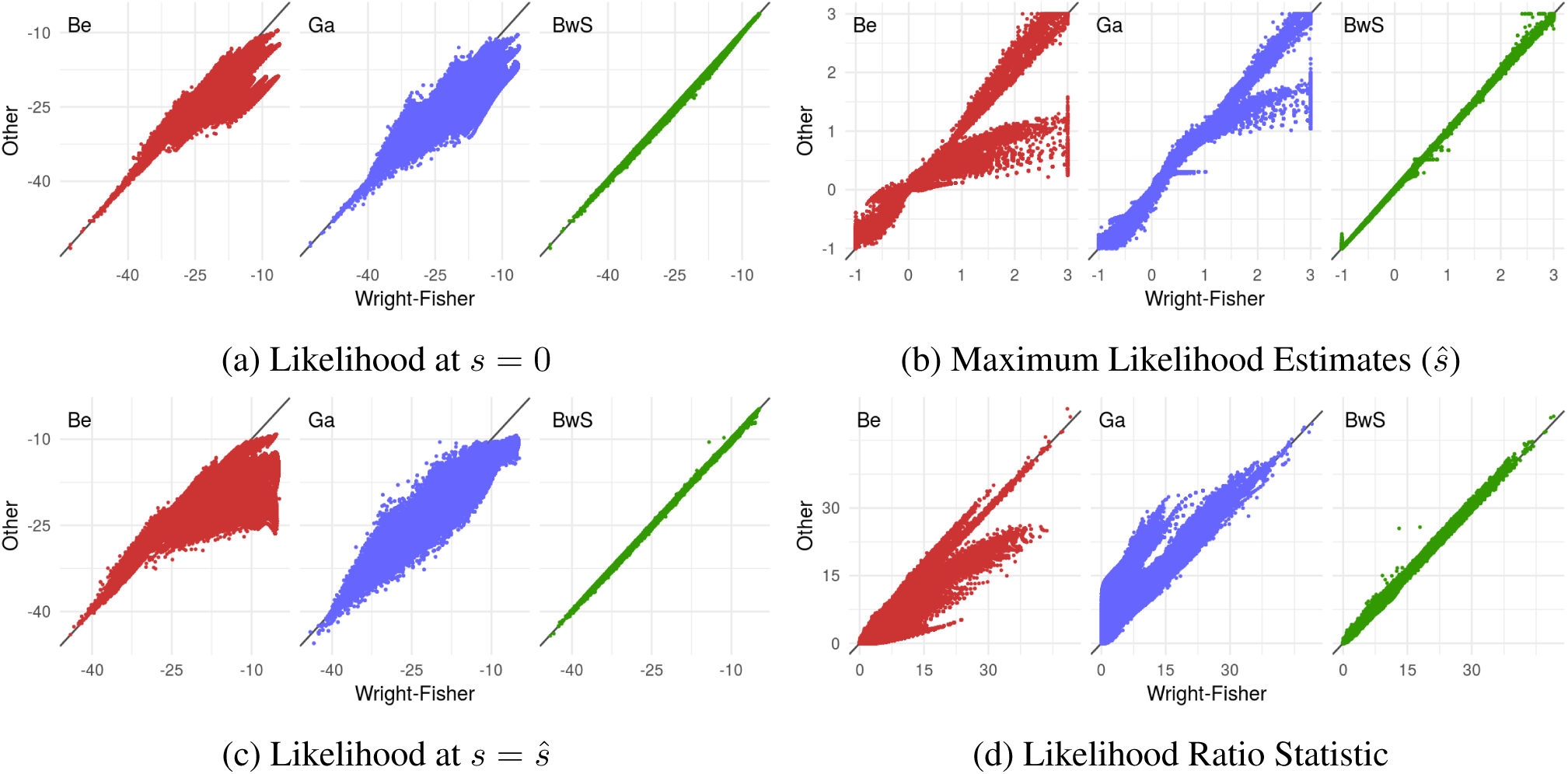
Comparison of three continuous approximations to the WF transition for likelihood-based statistics. In each subplot, Be, Ga, BwS indicate the compared model among Beta model, Gaussian model and Beta with spikes model

One reason why the BwS model is a better approximation to the WF than the Beta and the Gaussian is that it includes fixation probabilities. In the HMM framework, fixation events are considered by this model while others fail to correctly interprete vanishing observed allele frequencies. Indeed, when fixation events are unlikelily, all models approiximate efficiently the Wright-Fisher and the likelihood approximation is good. Fixation events depend mainly on the total duration of sampling *T* (see Figures S4, S5, S6, S7). For large values (*T/N*_*e*_ > 0.5) early fixations events (apart from the pathological cases mentionned above) are not interpreted by the Beta and Gaussian models as fixations. These models will consider that the constant trajectory (only 0 or 1 observed allele frequency since the fixation event) are consistent with a neutral process. The resulting maximum likelihood estimator therefore underestimates the true *s* (Figure 4b). As the maximum is not well found by Beta and Gaussian (Figure S8), the likelihood ratio statistic is inconsistent and is not calibrated as a *χ*^2^(1) distribution.

### Inferring selection using the Beta-with-spikes model

Given that the beta with spikes model closely mimics the Wright-Fisher model in the HMM framework, we studied the statistical properties of the HMM with BwS transitions for population sizes of 1,000 and 10,000 haploids, which are computationally prohibitive when using the WF transitions. In terms of statistical properties, we studied the ability to detect selective loci and the accuracy in estimating the intensity of selection.

#### Detection of selection

The HMM framework developed here allows to derive a likelihood ratio statistic that can be used to test for the null hypothesis *s* = 0. Asymptotically, this statistic follows a *χ*^2^(1) distribution. However, the asymptotics in the HMM framework are measured in terms of the number of dates sampled. In our simulations, and most likely in typical real datasets, this number is not very high (10 here). Our first goal was thus to evaluate whether the *χ*^2^(1) was a good fit to statistics computed under the null hypothesis. Then we evaluated its power under different alternatives.

#### Calibration under the null

We checked for the good calibration of the BwS under neutrality. Figure 5 shows on the y axis the quantiles of the empirical likelihood ratio distribution and on the x axis, the corresponding quantiles of a *χ*^2^(1) distribution. Generally the fit of the LR to the theoretical distribution was good although the test would be slightly conservative for large *T/N*.

**Figure 5:**
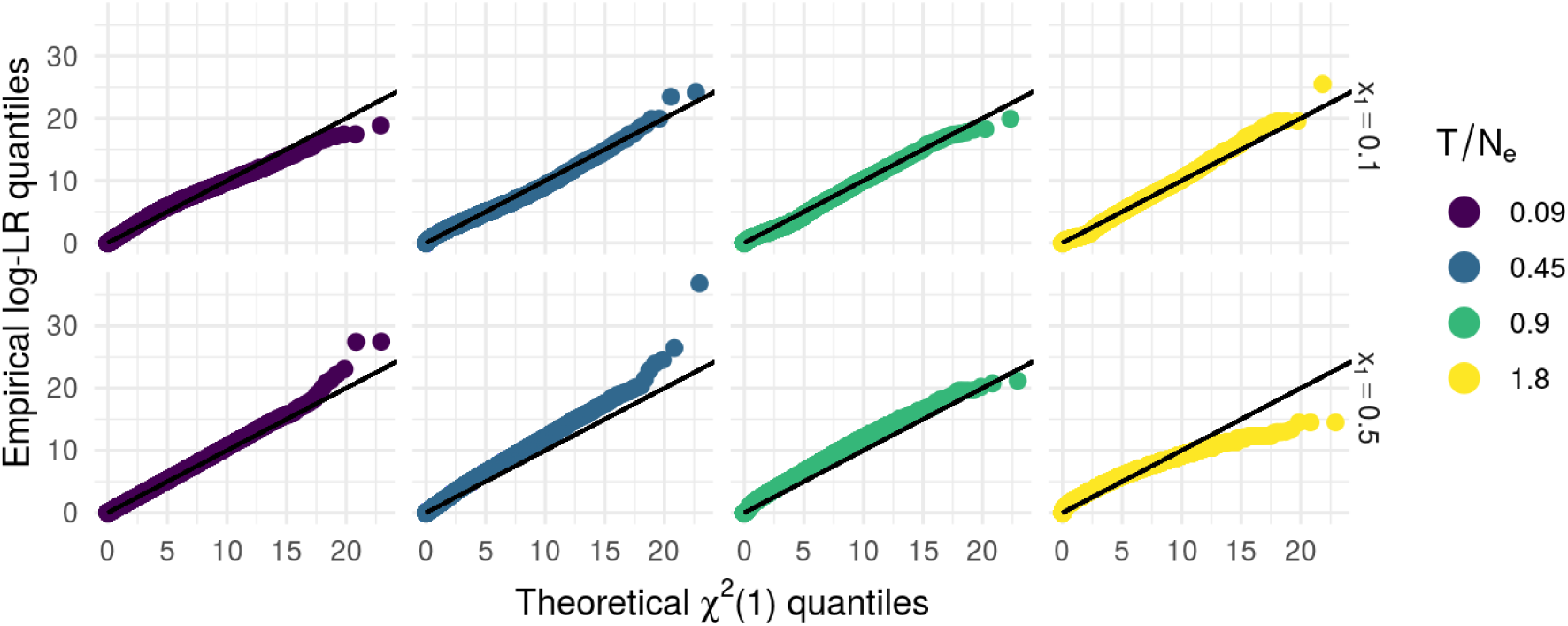
Calibration of *λ*(*y*) under the null: comparison of empirical vs theoretical *χ*^2^(1) quantiles for different population sizes, initial allele frequencies and inter-sample times.

#### Power

With the LR statistic and the *χ*^2^ quantiles, the empirical power of the test can be computed for different scenarios. As in this case we know that all true *s* parameters are non negative, we considered that all negative ŝ having a high LR should not be considered as a success in detecting selection. Indeed, even if the LR is high in this case, the final conclusion of such results would be that the wrong allele is under selection which is even worse than not detecting selection.

Figure 6 shows the power of the test as a function of *N*_*e*_*s* (Figure 6a) and as a function of *T/N*_*e*_ (Figure 6b), for two different initial allele frequencies 0.1 and 0.5. As expected, the power increases with *N*_*e*_*s* (Figure 6a). We can see also that for larger *T/N*_*e*_, the power starts to increase for smaller *N*_*e*_*s*. This can be explained by the fact that *T* is larger in such scenarios so selection has more time to affect the allele frequency and even small *N*_*e*_*s* give rise to trajectories that are consistent with selection. However, when starting at an initial frequency *x*_1_ = 0.5 the power decreases when *T/N*_*e*_ is too high. In these cases, the probability to observe large variations of allele frequency under neutrality becomes larger and so the ability of the LRT to distinguish neutrality from selection decreases.

**Figure 6:**
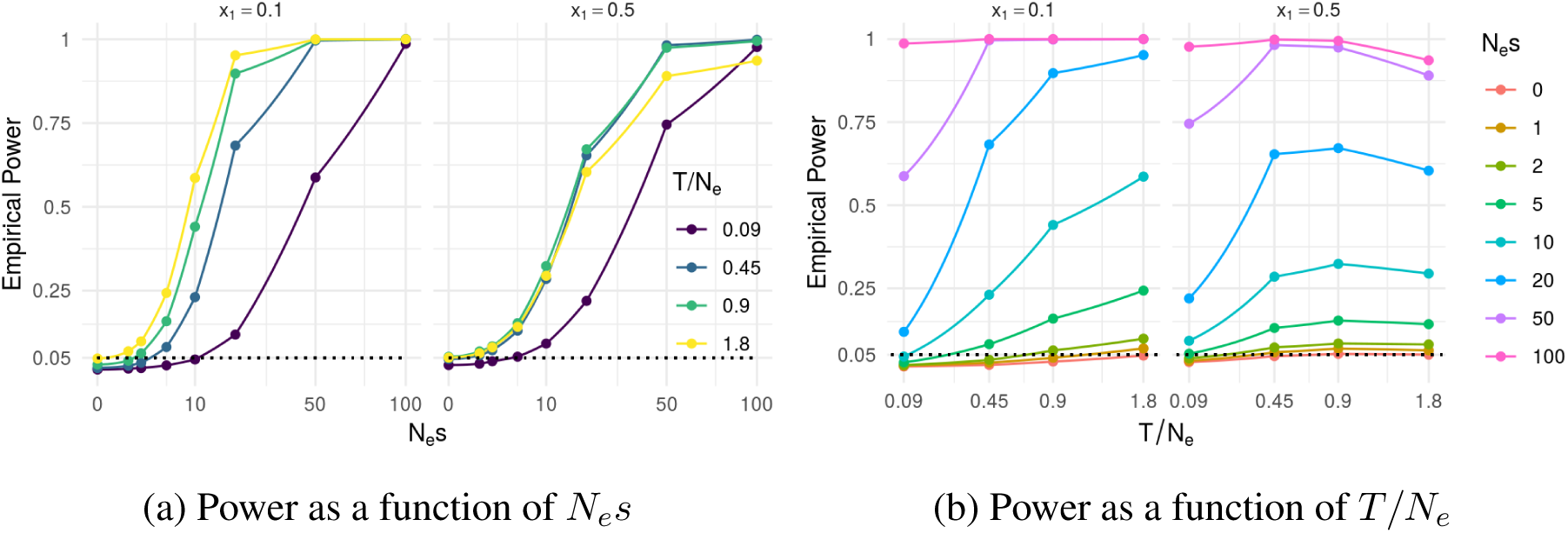
Power of the likelihood ratio test under different simulated scenarios.

#### Parameter Estimation

The Figure S9 shows that under parameters considered, *N*_*e*_ has no influence on estimation, except for large *N*_*e*_*s*, for which the maximum likelihood estimator has a higher variance for small *N*_*e*_. In this case, the diffusion approximation is not valid so different *N*_*e*_ are not equivalent for given *N*_*e*_*s* and *T/N*_*e*_.

Figure 7 shows the MLE distribution for different *N*_*e*_*s* and *T/N*_*e*_ pooling results from different *N*_*e*_. It shows that the estimation is more accurate (with lower variance) for large values of *T/N*_*e*_. Moreover, when fixation events (*T/N*_*e*_ *<* 0.5) are unlikeliy, the estimation is unbiased. On the other hand, when fixation event are likely (*T/N*_*e*_ > 0.5 and *N*_*e*_*s* > 50), the trajectories that are not filtered out are those that are consistent with lower values than the true *s* and therefore the MLE underestimates the true *s*.

**Figure 7:**
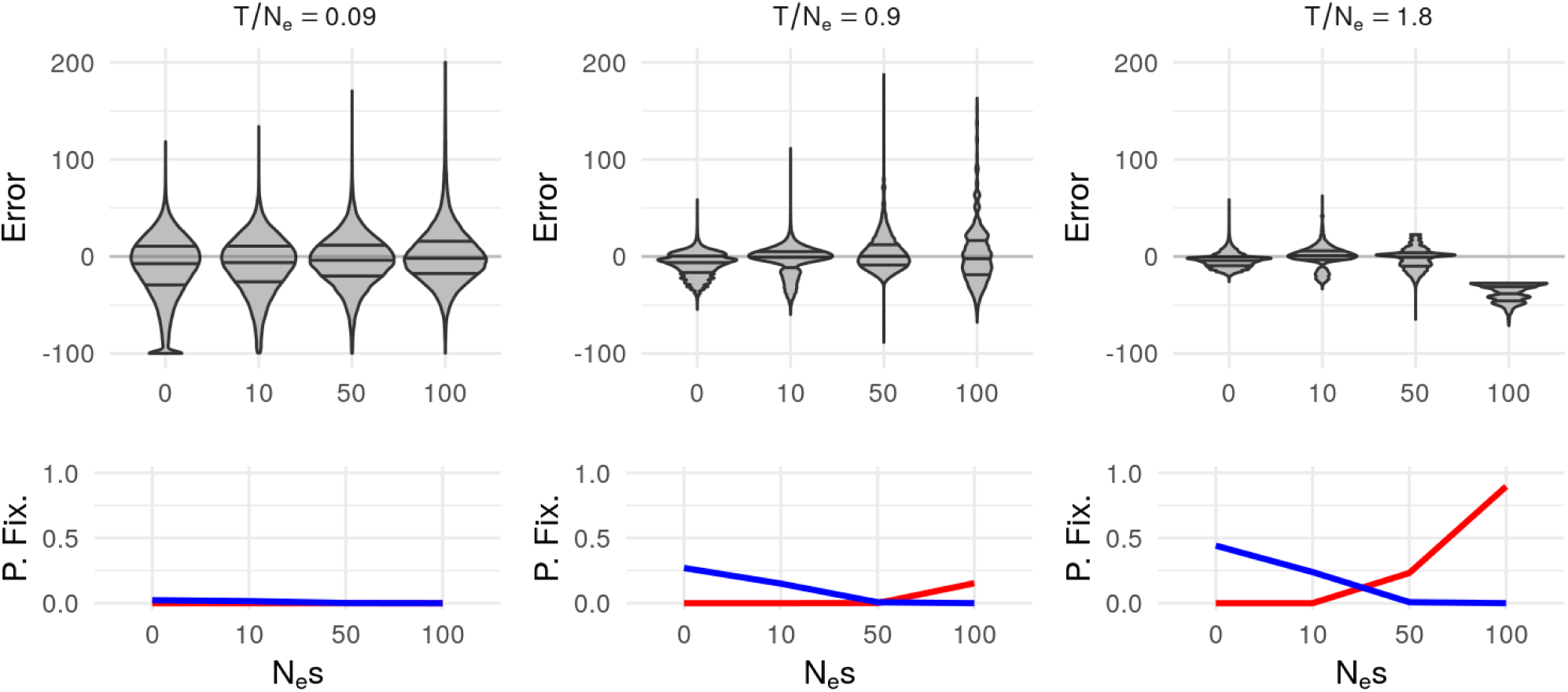
Estimation error distribution in different scenarios for *x*_1_ = 0.1. Each column stands for a scaled time parameter *T/N*_*e*_ ∈ {0.09, 0.9, 1.8 }. The first line indicates the absolute error distribution in each scenario and the second line represents in each cases, the proportion of rejected trajectories (red lines for fixations in 1 and blue lines for fixations in 0)

### Analysis of a selection experiment

To illustrate the statistical performance of the HMM approach on a real data set, we applied it to allele frequency trajectories observed at 40,199 SNPs in two lines of chicken divergently selected for the intra muscular ultimate pH (pHu). This dataset was previoulsy analyzed by Bihan-Duval et al. (2018) using the FLK Bonhomme et al. (2010) and hapFLK Fariello et al. (2013) tests: considering each generation of the experiment independently of the others, these authors looked for regions showing a significant excess of allele (for FLK) or haplotype (for hapFLK) frequency differentiation between pHu+, pHu- and the founding population. Thus, this case study offers the opportunity to compare the regions detected under selection by two different approaches: genetic differentiation and time series.

We first estimated population size in the two lines using the NB R package (Hui and Burt, 2015) and found *N*_*e*_ = 157 haploids in pHu+ and *N*_*e*_ = 123 haploids in pHu-. Based on these values, we applied the HMM in each line and detected a total of 39 significant SNPs under selection at a FDR of 5% (Table S1). These SNPs were clustered in 12 genomic regions (Table 2, Figure S10): eight of them were detected only in the phu+ line, one was detected only in the phuline and three other were detected in both lines. Moreover, for almost all significant SNPs, the allele that was selected in one line (positive value of ŝ) was also counter selected in the other line (negative value of ŝ) (Figure 8). This is consistent with the experimental design of divergent selection between the two lines.

**Table 2:**
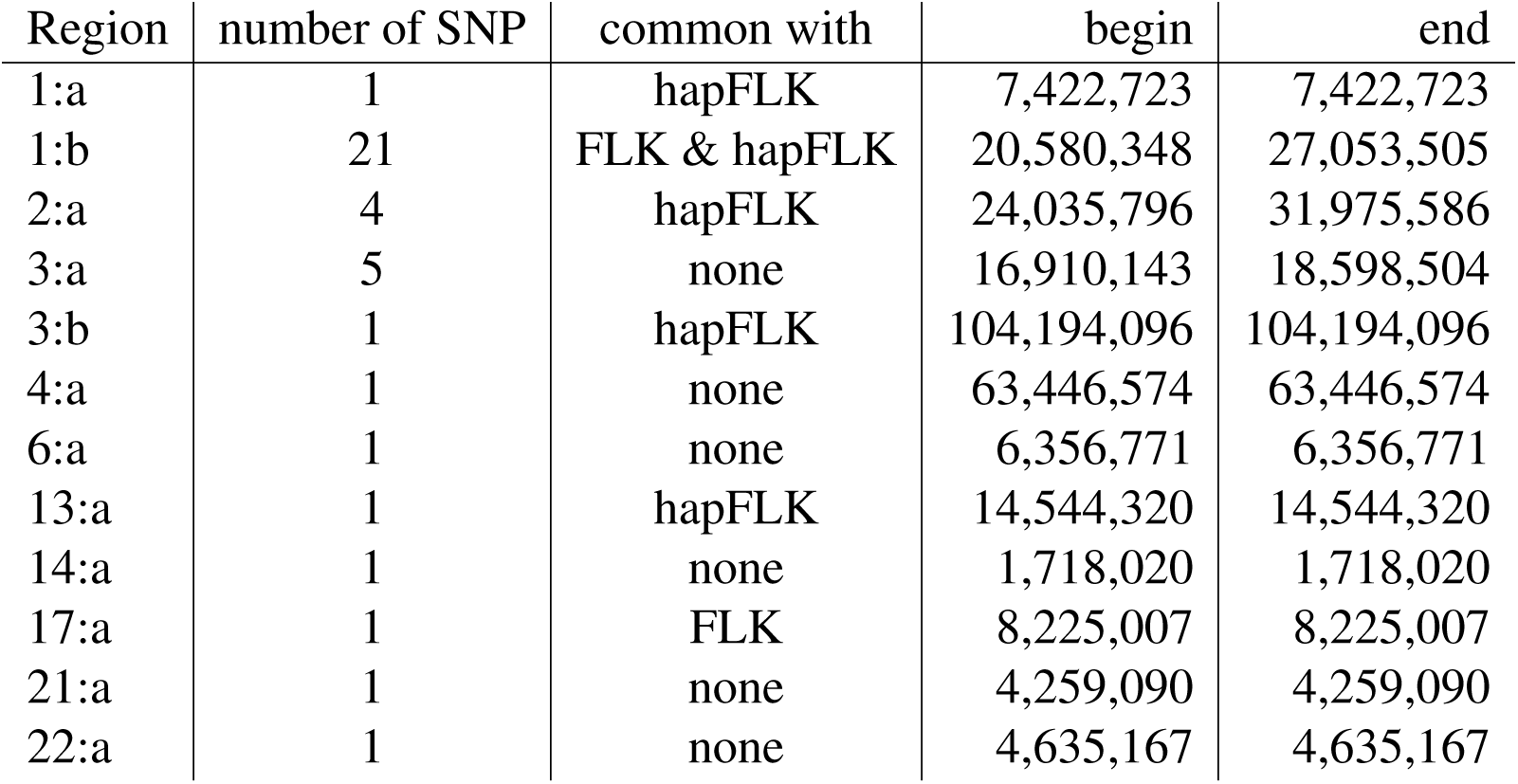
Significant regions (for HMM)

**Figure 8:**
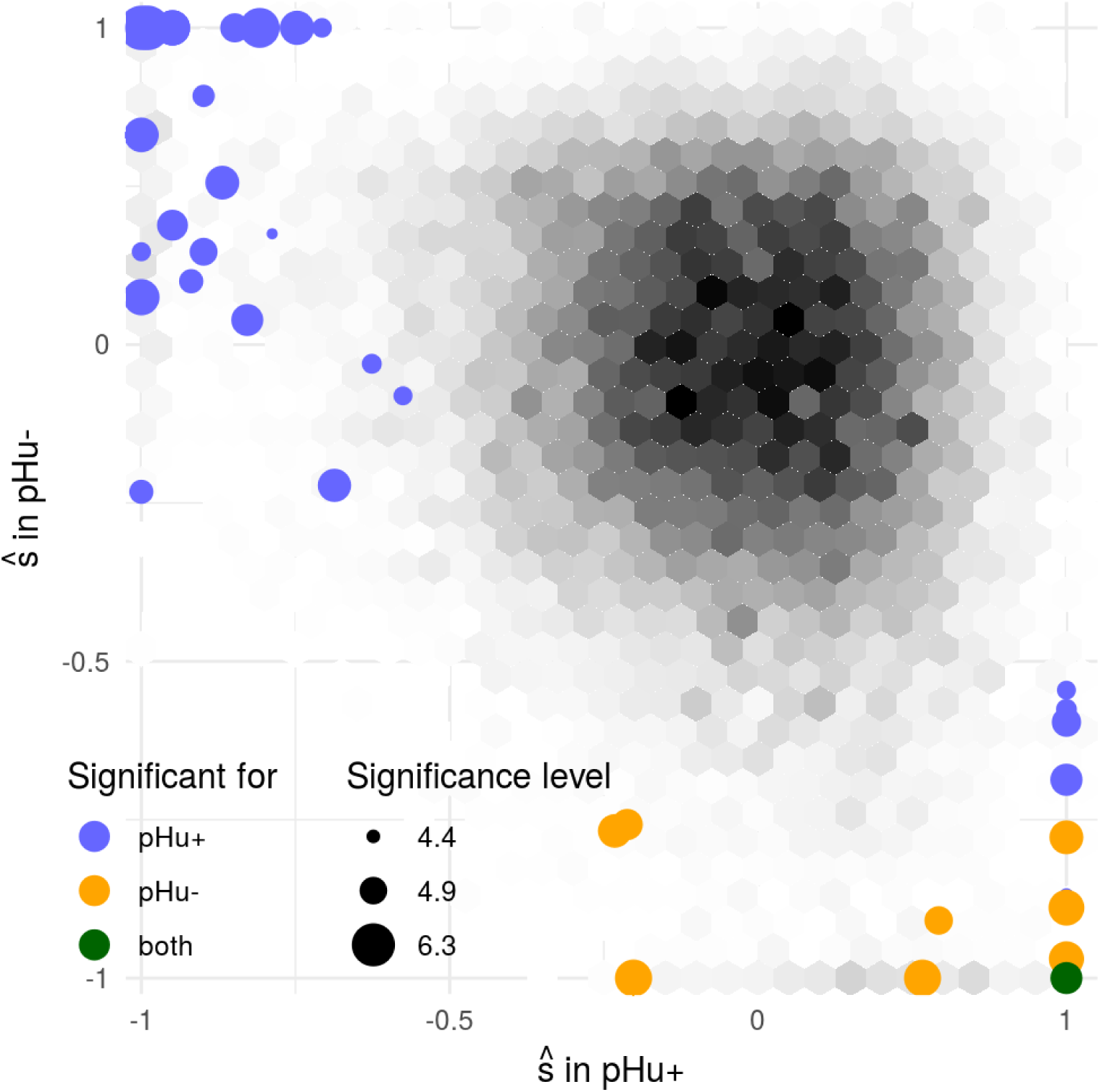
Distribution of ŝ in pHu-vs pHu+. Levels of grey represent the density of ŝ, significant SNP are represented in colors, the size representing the *-* log_10_ associated p-value

Among the 12 regions detected by the time series approach, six were detected by FLK or (and) hapFLK and six were new. SNPs in these six new regions were associated to an allele frequency significantly increasing or decreasing in one line but rather constant in the other line (Figure S11): in such cases, genetic differentiation was elevated but did not result in a significant FLK or hapFLK p-value. In contrast, SNPs detected only with FLK were generally associated to a moderate allele frequency change in both lines, that was not found significant when analyzing each of them independently (Figure S12). Overall, the differentiation and time series approaches were thus found complementary. Note also that the number of regions detected (i. e. the detection power) with the time series approach was very similar to that obtained by FLK. Detection power was significantly higher with hapFLK, which was related to the use of haplotype information.

## Discussion

In this study we have described a general statistical framework for the analysis of genetic time series. It is built upon combining a Hidden Markov Modeling approach, first proposed by Bollback et al. (2008) in the context of genetic time series and parametric approximations to the Wright Fisher process. Within this framework we have shown that using the Beta-with-spikes distribution proposed by Tataru et al. (2015) to approximate the Wright Fisher transitions provided a statistical inference comparable to that of the Wright-Fisher model while having a computational cost that does not depend on population size. We have shown this framework allows to perform hypothesis testing by relying on the asymptotic distribution of the Likelihood Ratio Statistic and that the maximum likelihood estimate of the selection intensity parameter had generally good statistical properties.

### Advantages and limits of parametric approximations to the Wright Fisher process

Using parametric distributions to approximate the Wright-Fisher transitions requires to know or approximate the moments of the Wright-Fisher process, including fixation probabilities in the case of the BwS model. We have shown, in the lines of previous work by Lacerda and Seoighe (2014); Terhorst et al. (2015) that this could be done using a Taylor expansion of the fitness function (see Supplemental File S1). Despite generally good performance of the approximation of moments, there still exists situations were the approximation performed poorly. For the BwS model, this only happened for low starting allele frequencies (*x*_*s*_ ≲0.1), large number of generations (*t/N*_*e*_≳0.1) and strong selection. Note that the poor performance for low starting allele frequencies is relevant for all *underlying true* allele frequency as for computing the likelihood the transition is integrated over all possible starting frequencies in the forward algorithm. So this poor performance could have impacted our simulation results. However, we showed that the WF likelihood was still well approximated by the BwS model. In cases where better approximation of the WF moments might be required, different approaches could be followed. For small population sizes (*N*_*e*_≳1000), the WF moments can actually be computed exactly using equation (4) but this computation quickly becomes computationally prohibitive (the cost is approximately 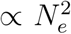). A solution would be to develop the Taylor expansion some degrees further and possibly include correction terms as was done by Terhorst et al. (2015). This could be important for datasets covering large timescales such as in ancient DNA studies where sampling times can be widely spread.

Here, we compared three different parametric distributions for approximating the Wright-Fisher transitions. The Gaussian distribution is a widely used approximation to the evolution of allele frequencies, starting from Nicholson et al. (2002), including for the analysis of genetic time series (Lacerda and Seoighe, 2014; Terhorst et al., 2015). Our results show that this approximation is only valid for small time scales (in units of *t/N*_*e*_) and when the starting minor allele frequency is high. The Beta distribution has been less used in this context (Gompert, 2016; Siren et al., 2011) although we have shown that it offers a better approximation to the transitions than the Gaussian model. However these two models present poor performance when it comes to approximating the likelihood of data simulated under the Wright-Fisher model. This it not true for the Beta-with-spikes model that provides good approximations both for the transition kernel and the likelihood. This is most likely due to the fact that it incorporates fixation probabilities that become large when the other models fail (large selection intensity, large intersample time, low minor allele frequency).

The BwS model should therefore be preferred to the other two at least in the context of analyzing genetic time series using Hidden Markov Models. Given the good performance of the BwS model in this context, it could be an interesting model to use in more general contexts. One interesting extension would be to use the BwS in the context of time series data sampled in multiple populations. However, one important limitation of using the Beta distribution (with or without spikes) for such data is that the multivariate Beta distribution is essentially unusable in practice. Using this class of distribution in a multiple populations context therefore requires developing a specific factorization for calculating the multivariate likelihood. An example of such an extension for the problem of estimating branch length in population trees can be found in Tataru et al. (2015). For such data, the Gaussian model remains much more practical but still suffers from the limitations mentioned above.

In this study, we only focused on comparing parametric distributions to approximate the WF transitions. Another standard approximation to the Wright-Fisher process under selection that was not considered here is the diffusion process. This process has been shown to provide a good approximation to the WF process. However this approximation is only valid for large *N*_*e*_ and small *N*_*e*_*s* values and has two additional important limitations. The main limitation is that using the diffusion model in practice implies solving the diffusion equation which is a partial differential equation. One approach to do so is to write the transition distribution as an infinite sum of elementary solutions of the diffusion equation (Song and Steinrücken, 2012). This approach was considered in a HMM for genetic time series by Steinrücken et al. (2014) but exhibited prohibitive computational cost. Another approach is to consider numerical schemes to approximate the transition density (Bollback et al., 2008; Zhao et al., 2013). These approaches involve numerical integrations and are also computationally demanding. The second limitation of the diffusion process is that it does not naturally incorporate the fixation probabilities at 0 or 1. However, Zhao et al. (2013) showed that the discretized diffusion transition density had atoms on these edges that could be used to represent the fixation probabilities. This suggest that the diffusion approximation could be extended to include fixation probabilities. This would require specific developments and evaluation.

Overall, the choice of a particular transition model for genetic time series should depend on the study to analyze. For small *N*_*e*_, WF calculations are tractable and should be preferred. For larger *N*_*e*_, the diffusion approximation could be improved but parametric approximations seem more adapted (Lacerda and Seoighe, 2014). In the class of parametric distributions, our results show that the Beta-with-spikes model is probably more flexible as it is adapted to both large *N*_*e*_ and a wider range of *N*_*e*_*s*.

### Inference of selection from genetic time series

In principle, genetic time series can provide a lot of information on selection at a locus. This comes from the fact that the allele frequency trajectory underlying the observed sample is greatly affected by the selection process. Indeed we found that in many scenarios that we simulated likelihood inference provided essentially unbiased estimates of the selection parameter. The power to detect selection was substantial for a wide range of situations when the selection intensity *N*_*e*_*s* was greater than 10. For large populations this corresponds to a relatively modest fitness effect. For example for a population of size 10,000 this corresponds to *s* = 0.01, *i.e.* a 1% increase in fitness for an individual homozygote for the beneficial allele compared to the other homozygote.

Our approach builds upon likelihood inference and in particular maximum likelihood estimation. While this approach was found very efficient in most cases, the maximum likelihood theory could not be applied to allele frequency trajectories where the reference allele was fixed or lost between the first and second sampling dates. Indeed, the best explaination for such events is ŝ = *-*1 (for allele loss) or ŝ = ∞ (for allele fixation), in other words the MLE is reached on an edge of the domain. The occurence of such events implies that the LR distribution under neutrality is not *χ*^2^(1). An approach to ensure properly distributed p-values in this situation would rely on performing neutral simulations to obtain empirically the distribution of the LR under the null. However, a more fundamental issue is that the ML estimator gives no accurate information about the true *s* here, because sampling dates are too distant to capture how fast the allele frequency increases (or decreases).

We chose to treat these trajectories specifically and exclude them from further analysis. However, the exclusion of fixed trajectories leads to an underestimation of *s*. This happens because the trajectories that are not excluded correspond to a biased sample of trajectories (those that reached fixation more slowly) and the resulting ŝ is lower than the true *s*. A way to compensate this effect would be to consider the transition process conditionnally on the allele not being fixed. This can be done in the method of moments framework by using the conditional moments, and with the diffusion model by modifying the diffusion equation (Ewens, 2004). However, this methodology may exclude a potentially large number of loci that can be properly analyzed within our framework, *i.e.* all trajectories including allele loss or fixation *later than* the second sampling date. In real data, the way to avoid such situations is through the experimental design by chosing an appropriate sampling strategy.

In this study we only examined one sampling strategy : 10 sampled dates equally spread along the total duration with 30 alleles sampled each time. With this strategy, we showed that the statistical quality of the model is driven by the *N*_*e*_*s* and *T/N*_*e*_ parameters: if the selection is too weak, increasing the duration considered is needed so as to capture its effect on the allele frequency trajectory. But if the total duration is too large, distinguishing the effects of genetic drift and selection is harder. The consequence of our results is that, for a given dataset, as the statistical accuracy of the estimation and the power of the test depend on the sampling dates distribution and the sample sizes, the interpretation of the results becomes more difficult. Some selected loci will escape detection although they can be potentially important for fitness. Hence, for a given dataset, we would suggest performing simulations (possibly using the scripts we used here) matched for sampling dates and sizes under different selection intensities to help interpreting selection scan results. Moreover, such simulations could actually be used to compare possible sampling strategies (*i.e.* depending on resources such as cost and sample availability) to optimize the power of the eventual statistical analysis. Deriving general guidelines for the optimization of sampling strategies in the context of genetic time series would require a dedicated study.

Our evaluation of the HMM framework focused on its performance to detect and estimate selection. To do so we have assumed that other parameters, in particular effective population size, were known. In addition, we assumed that each parameter was constant over the sampling duration. In practice most of these assumptions are deemed to be wrong. Regarding the estimation of *N*_*e*_, in our analysis of the real dataset, we used a two-step approach by first estimating *N*_*e*_ using Hui and Burt (2015) and then *s* at each locus. The issue with such an approach is that all loci are used to get the *N*_*e*_ estimate. If some of them are under selection, they will exhibit larger allele frequency deviation. The consequence is that the *N*_*e*_ estimate is going to be under estimated and it becomes harder to reject the null hypothesis (genetic drift). Hence, while this two-step approach is practical, it could be improved by jointly estimating *N*_*e*_ and *s*. The framework described here can in principle be used to do so as the HMM likelihood incorporates both *N*_*e*_ and *s*. This will require fitting the HMM model on multiple loci jointly as the information on *N*_*e*_ lies in the variance of the allele frequency trajectories over the whole genome. Also, it is now well established that populations of many species have experienced large changes in effective population time over time. There is also many situations where the selection intensity of an allele (expressed by *s* in our model) will change over time (*e.g.* time varying environmental conditions, selection for an optimum trait value …). In principle the joint model described above to jointly estimate *N*_*e*_ and *s* could also be extended to allow for parameters to vary in time (*e.g.* at each interval between sampling dates). The likelihood framework described here can form the basis of such models.

## Conclusion

Genome-wide assessment of the genetic diversity of a species is becoming more and more accessible in particular thanks to the development of cost effective genotyping and sequencing techniques. With the increase in genomic data, the evolution of genetic diversity in time becomes accessible either as a by-product of data accumulation or through dedicated projects in artificial populations (*e.g.* experimental evolution) or natural settings (*e.g.* ancient DNA sequencing). As genomic time series data become more widespread, their analysis requires dedicated methods that have good statistical and computational properties. In this study we have established a general statistical framework to this aim that we believe can contribute to further developments for the analysis of genetic time series.

---

**Figure S1:**
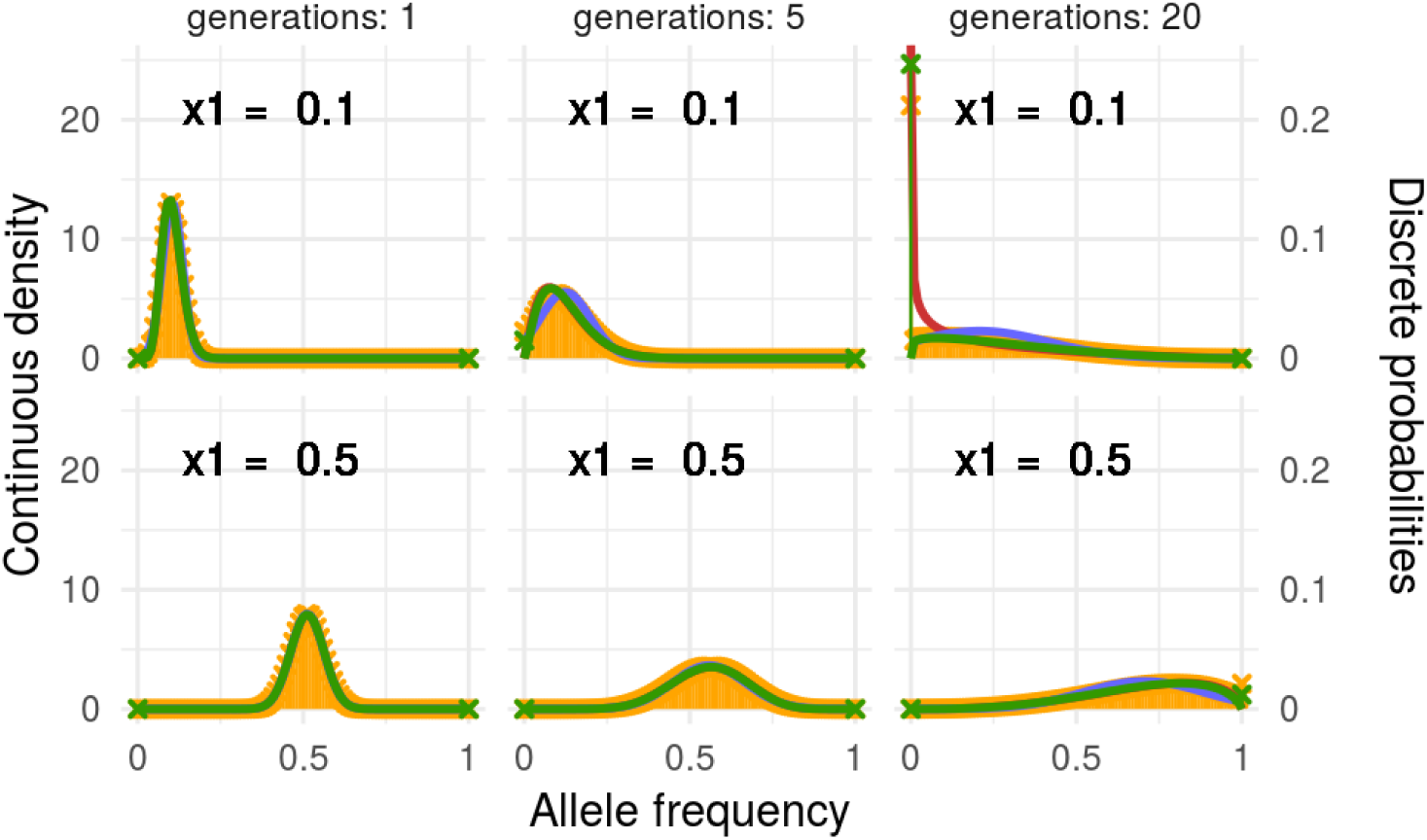
Some examples of approximations, for *N*_*e*_ = 100, *N*_*e*_*s* = 10, starting from a frequency of 0.1 or 0.5, during 1, 5 or 20 generations

## Supplementary Figures

**Figure S2:**
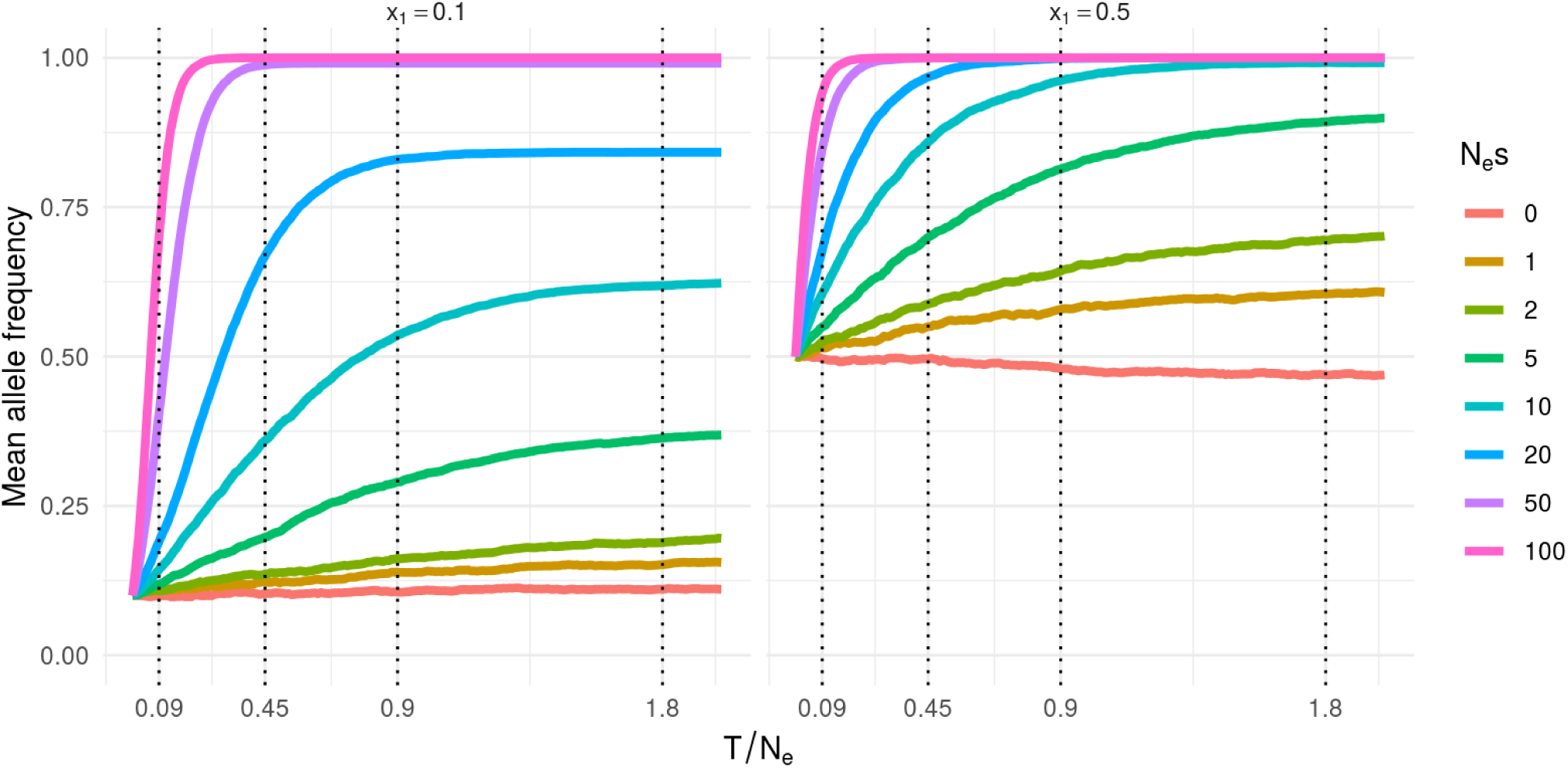
Mean allele frequency evolution.

**Figure S3:**
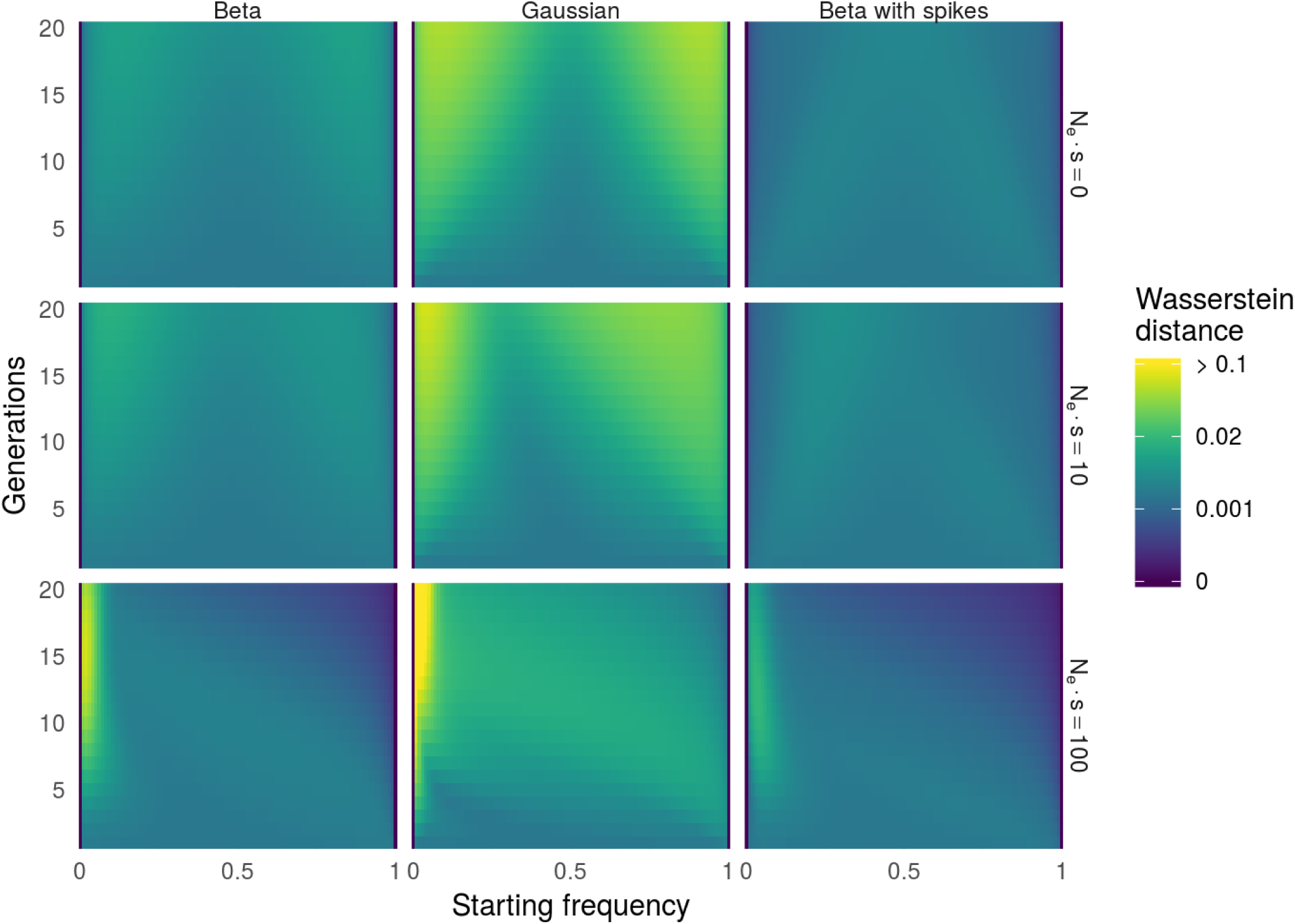
Wasserstein heatmap (true moments)

**Figure S4:**
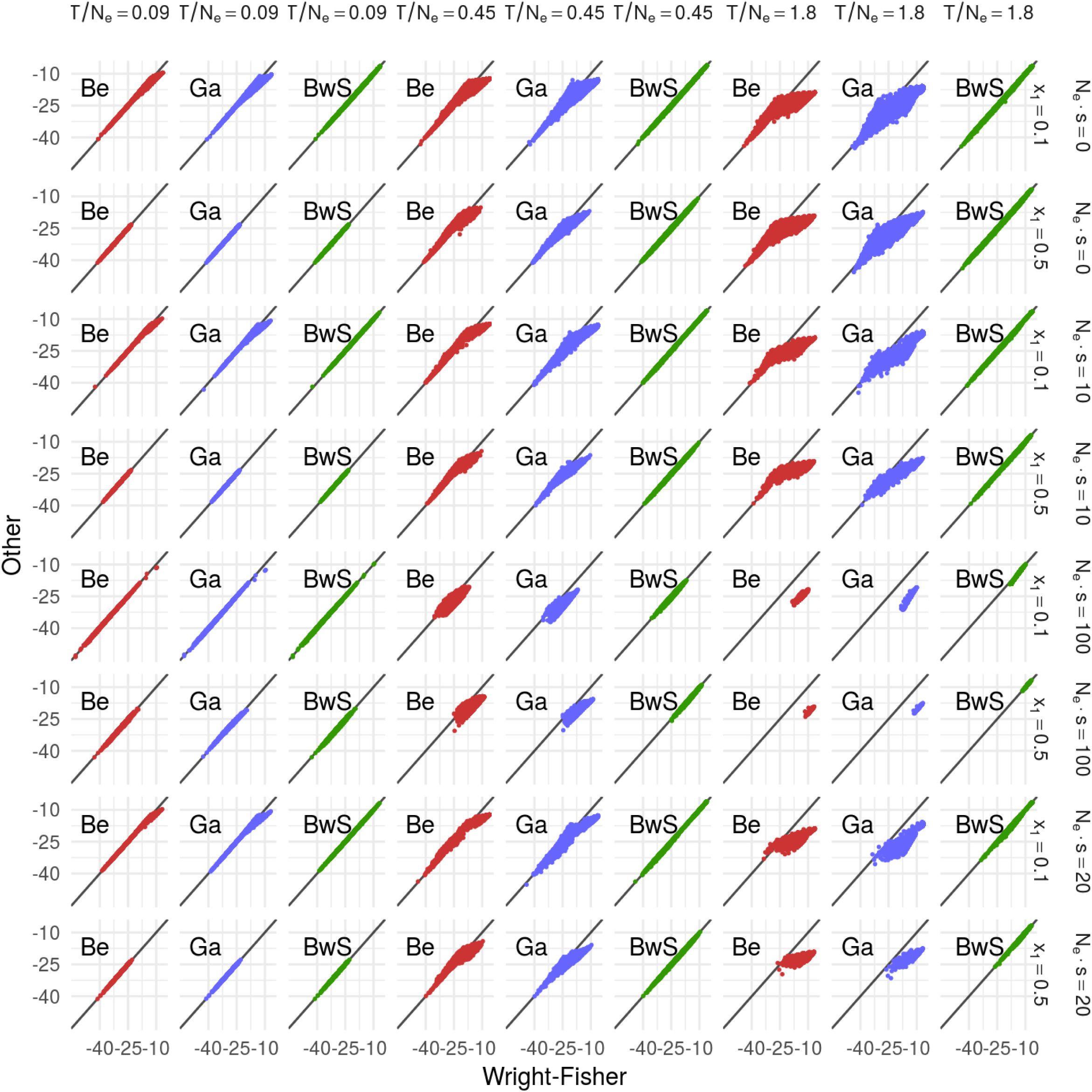
L0 comparison.

**Figure S5:**
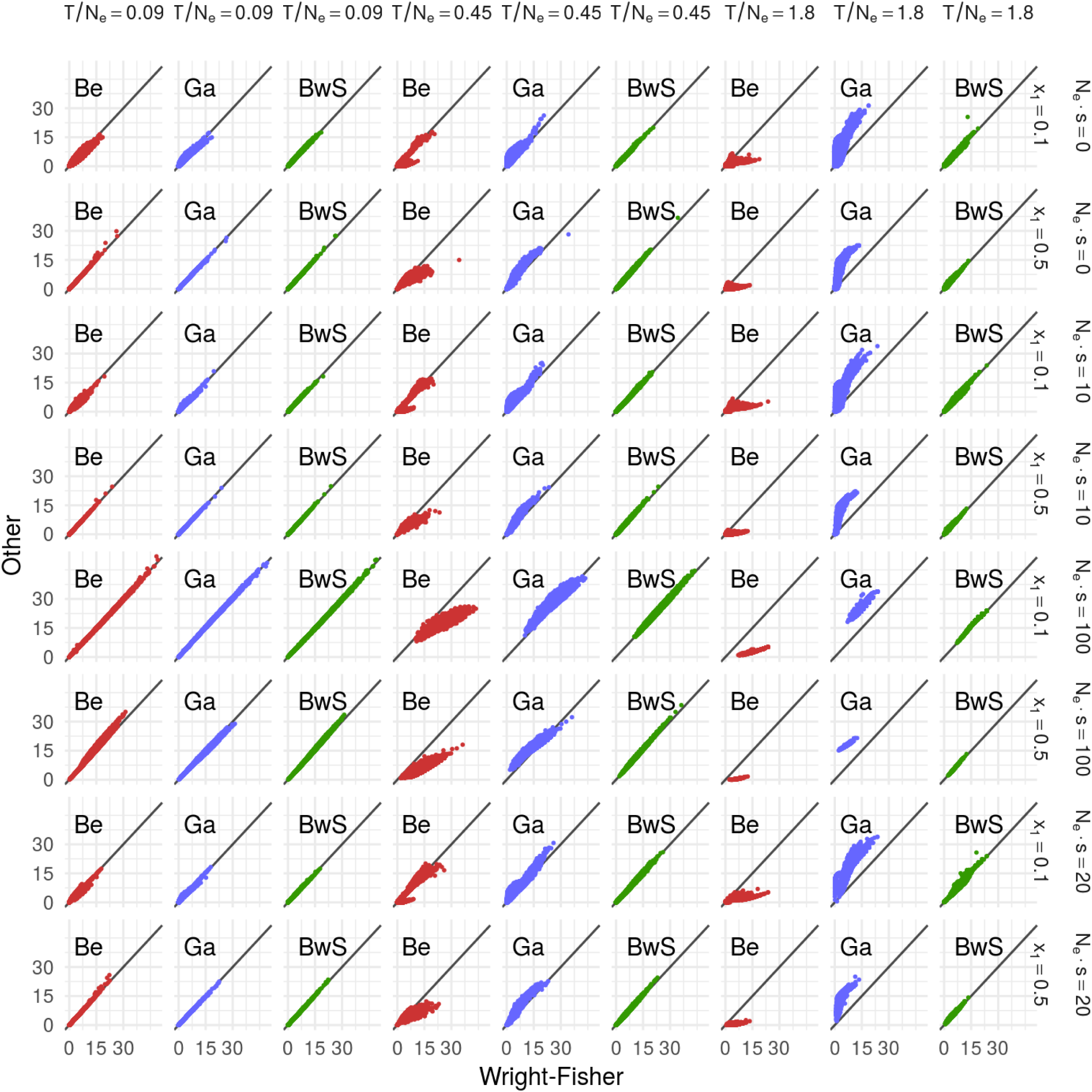
LR comparison.

**Figure S6:**
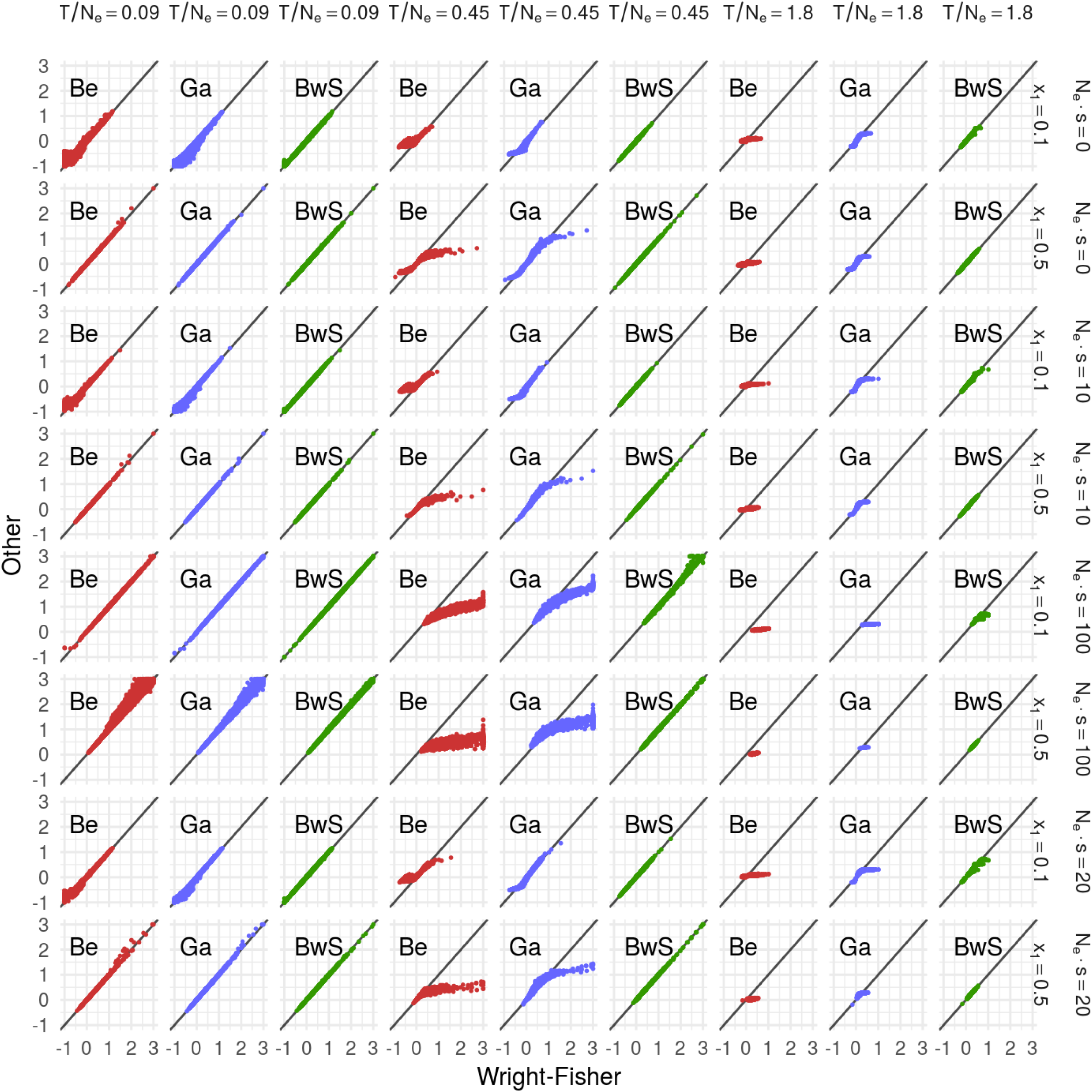
MLE comparison.

**Figure S7:**
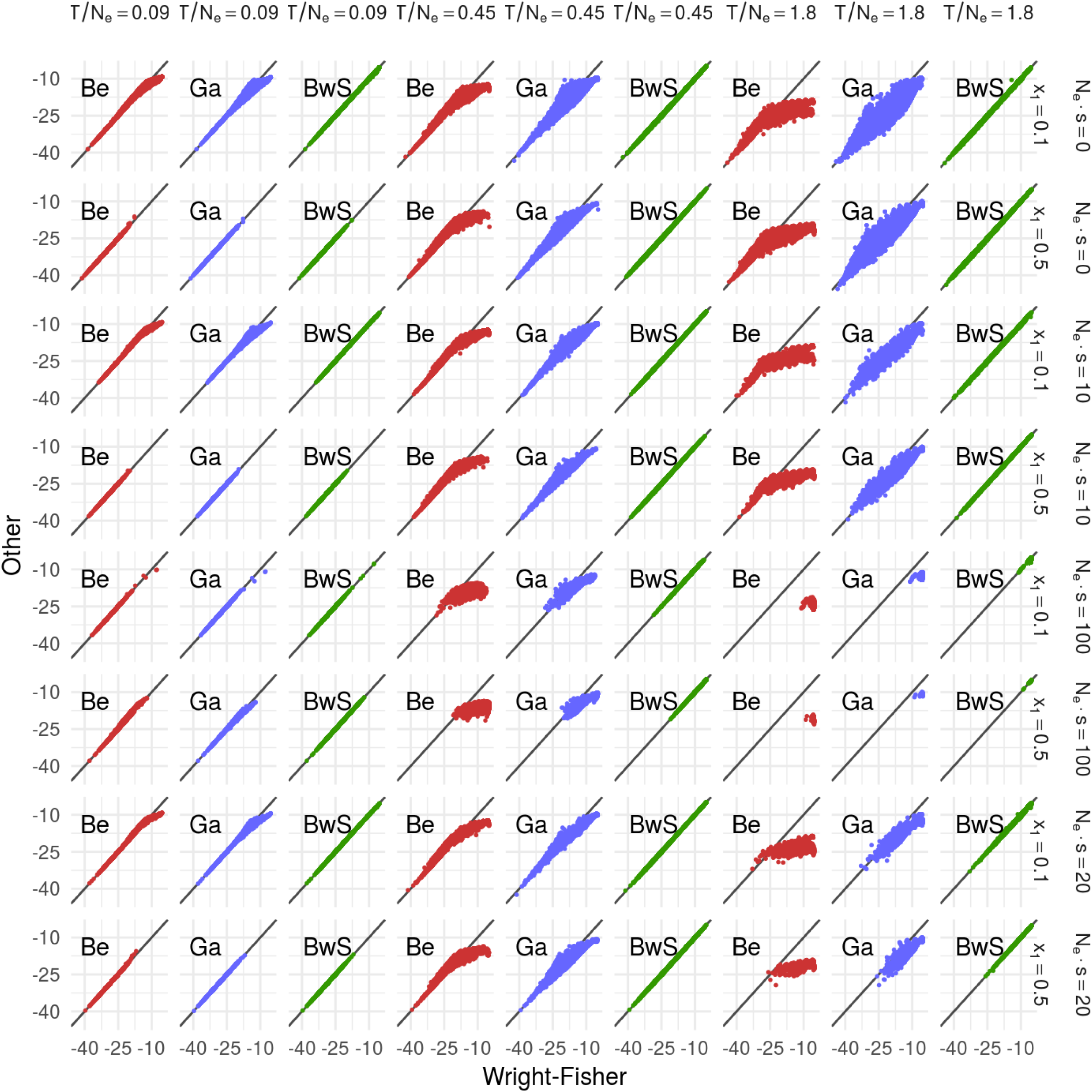
LMLE comparison.

**Figure S8:**
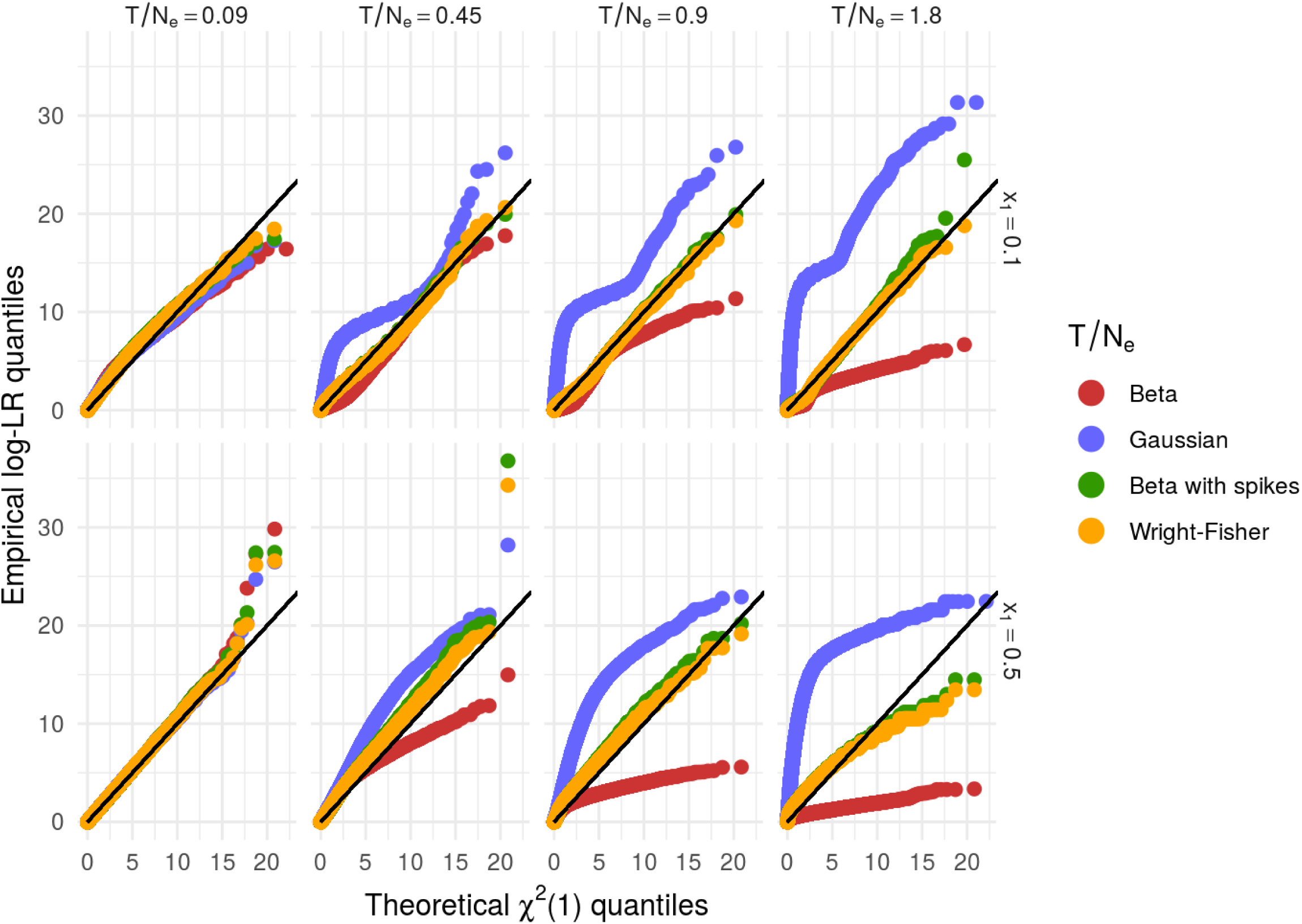
Calibration for other models.

**Figure S9:**
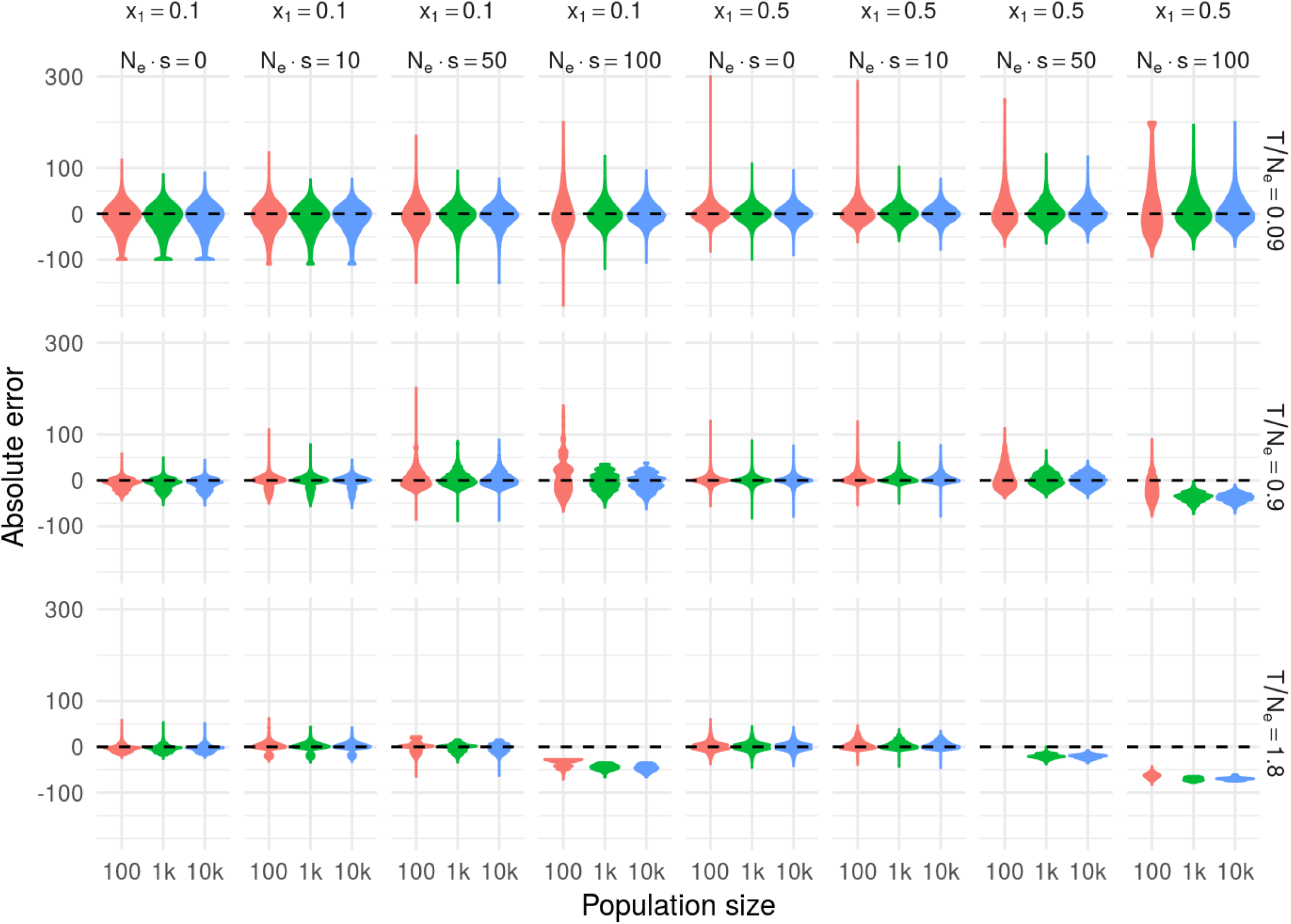
violins.

**Figure S10:**
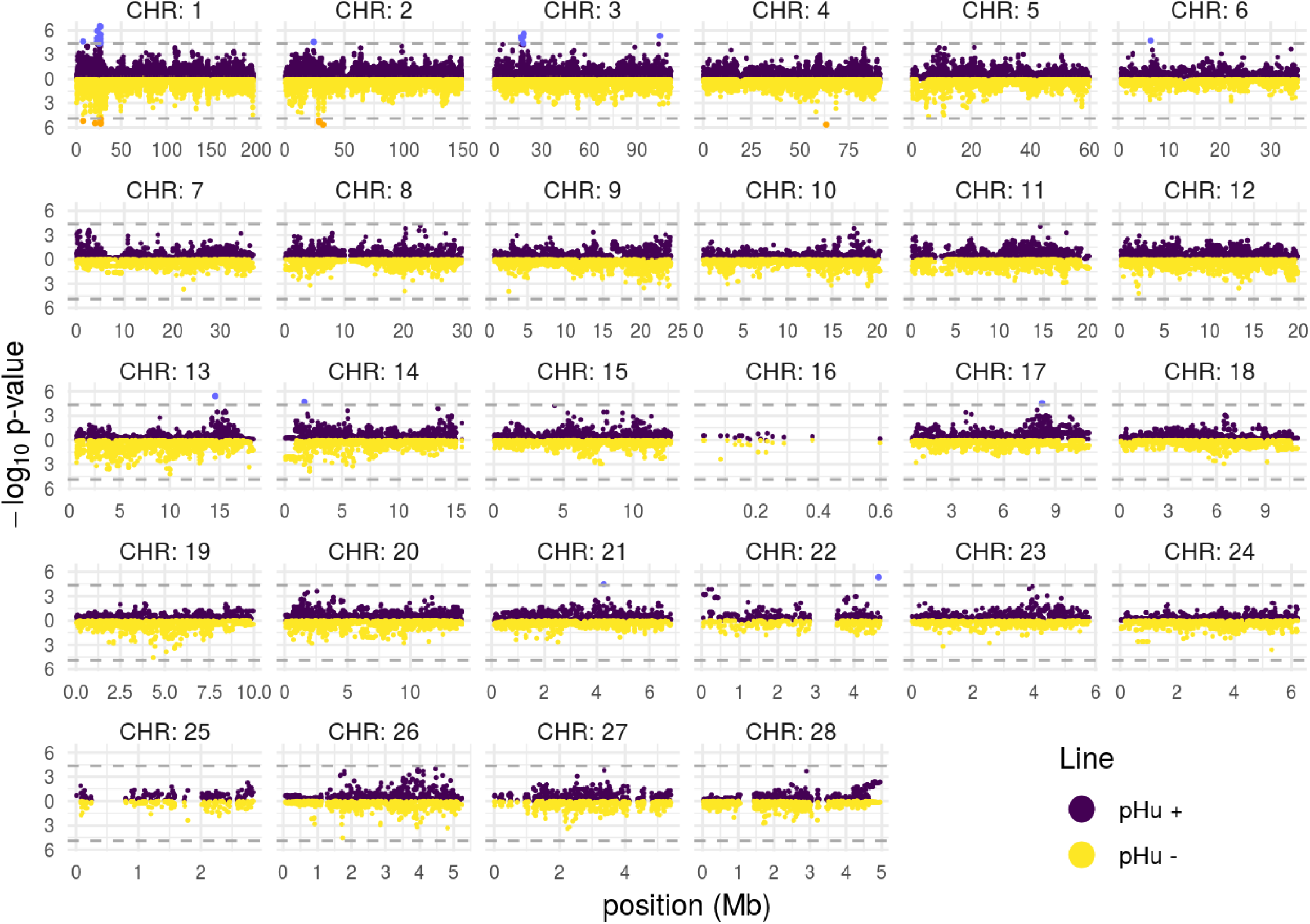
Manhattan plot.

**Figure S11:**
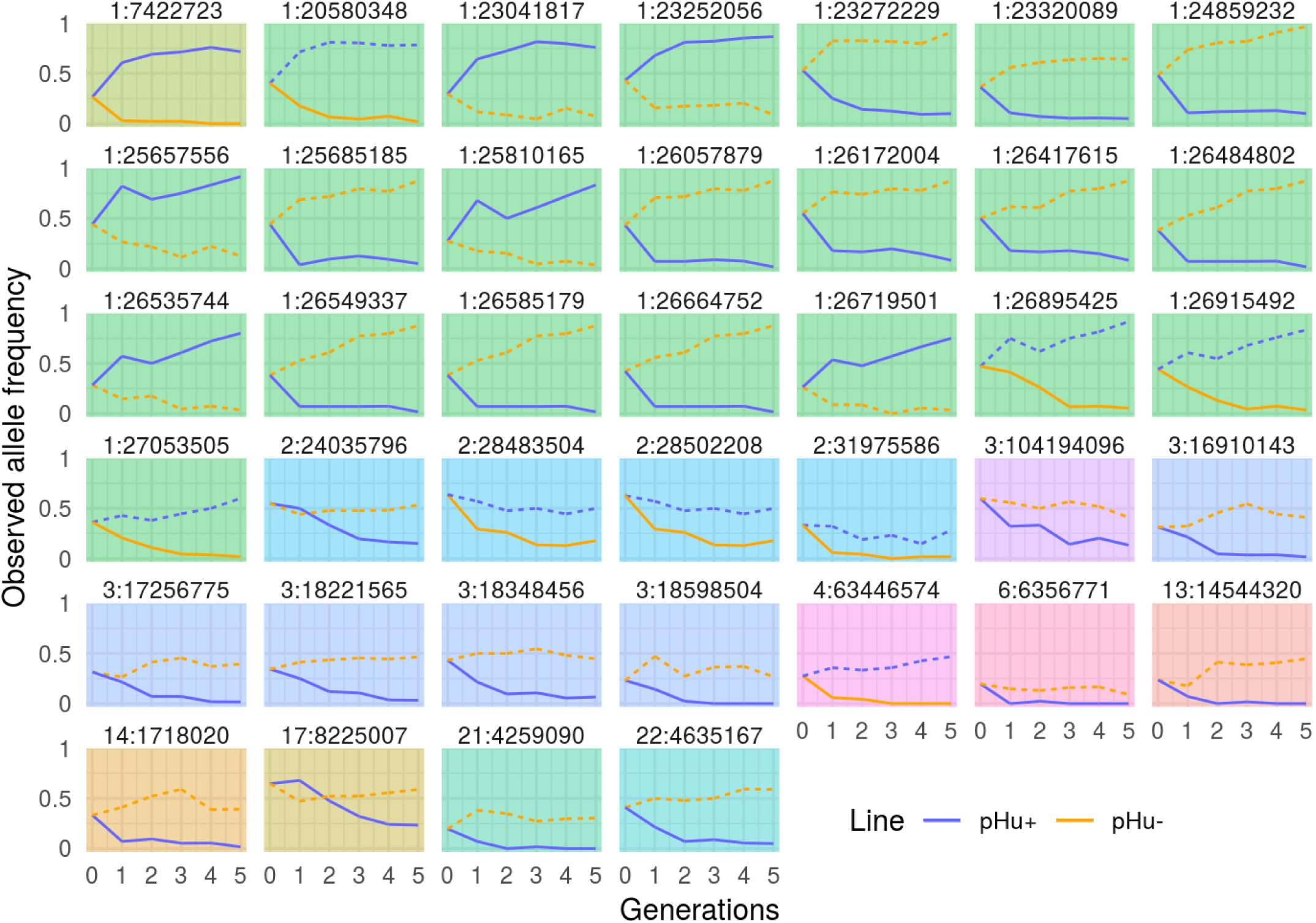
Allele frequency evolution of significants SNP.

**Figure S12:**
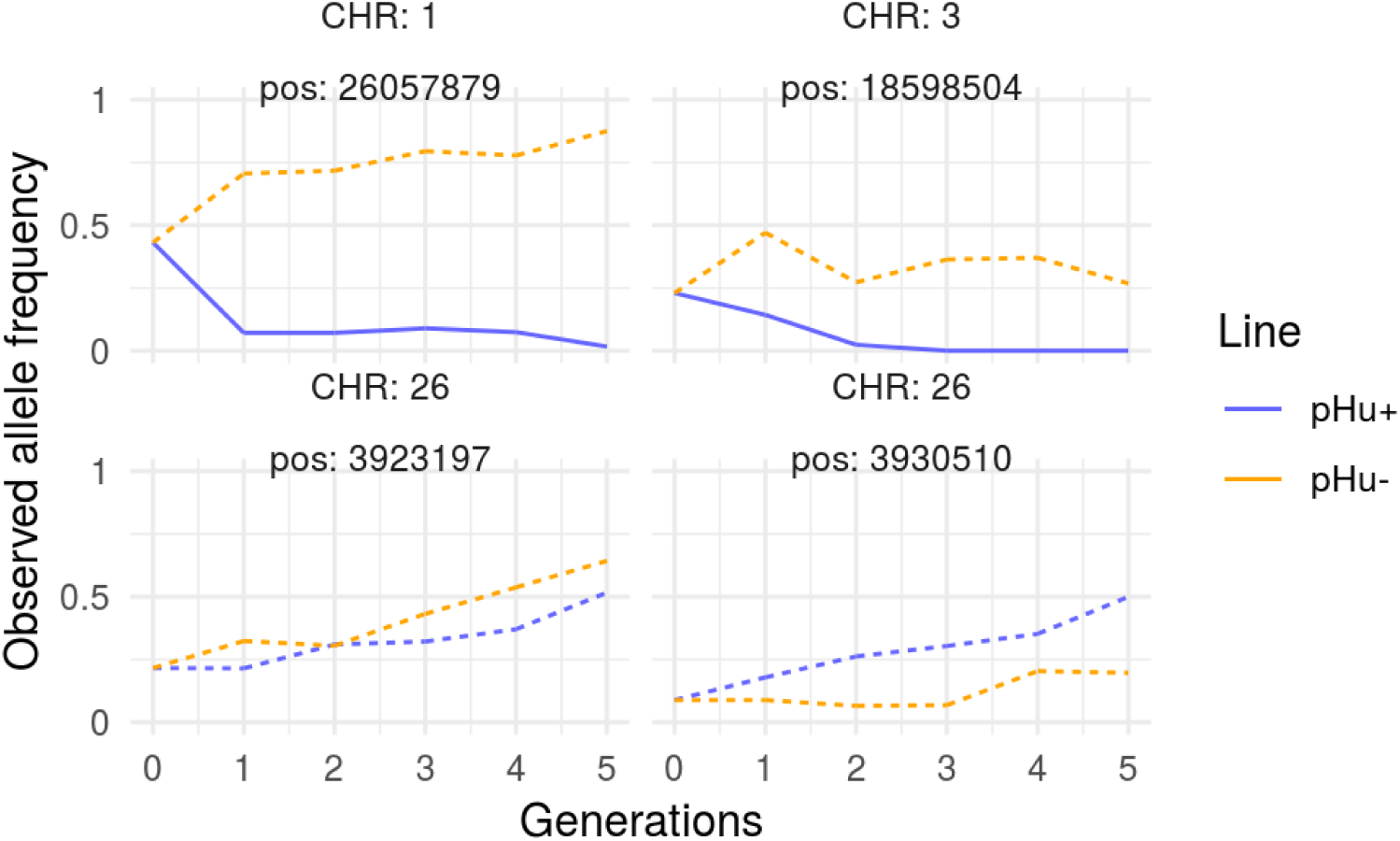
Allele frequency evolution of significants SNP.

## Supplementary Tables

**Table S1:** All Significant SNP with HMM approach.

| Region | CHR | position | $\log_{10}$ pval - | $\log_{10}$ pval + | Significant line |
| --- | --- | --- | --- | --- | --- |
| 1:a | 1 | 7422723 | 5.21077455 | 4.6074471 | both |
| 1:b | 1 | 20580348 | 5.47044894 | 3.5682896 | pHu- |
| 1:b | 1 | 23041817 | 1.75932274 | 5.0142726 | pHu+ |
| 1:b | 1 | 23252056 | 2.45288017 | 4.5683206 | pHu+ |
| 1:b | 1 | 23272229 | 2.83719381 | 5.9396324 | pHu+ |
| 1:b | 1 | 23320089 | 1.68565070 | 4.6247660 | pHu+ |
| 1:b | 1 | 24859232 | 4.60007599 | 4.3567040 | pHu+ |
| 1:b | 1 | 25657556 | 2.35853939 | 4.5047457 | pHu+ |
| 1:b | 1 | 25685185 | 3.44890614 | 5.0023590 | pHu+ |
| 1:b | 1 | 25810165 | 2.48970129 | 5.2272238 | pHu+ |
| 1:b | 1 | 26057879 | 3.56103427 | 6.4782438 | pHu+ |
| 1:b | 1 | 26172004 | 2.18477907 | 5.3908145 | pHu+ |
| 1:b | 1 | 26417615 | 2.96563878 | 4.5251675 | pHu+ |
| 1:b | 1 | 26484802 | 4.44639584 | 5.4721635 | pHu+ |
| 1:b | 1 | 26535744 | 2.56158176 | 4.9235733 | pHu+ |
| 1:b | 1 | 26549337 | 4.44639584 | 5.4721635 | pHu+ |
| 1:b | 1 | 26585179 | 4.44639584 | 5.4721635 | pHu+ |
| 1:b | 1 | 26664752 | 3.94536868 | 6.4587606 | pHu+ |
| 1:b | 1 | 26719501 | 2.92295423 | 4.3987421 | pHu+ |
| 1:b | 1 | 26895425 | 5.36602115 | 4.1493245 | pHu- |
| 1:b | 1 | 26915492 | 5.53265616 | 3.3652977 | pHu- |
| 1:b | 1 | 27053505 | 4.95463188 | 1.2118964 | pHu- |
| 2:a | 2 | 24035796 | 0.05527689 | 4.5356497 | pHu+ |
| 2:a | 2 | 28483504 | 5.33780455 | 0.7743481 | pHu- |
| 2:a | 2 | 28502208 | 5.14240939 | 0.7091620 | pHu- |
| 2:a | 2 | 31975586 | 5.65490823 | 0.5064980 | pHu- |
| 3:a | 3 | 16910143 | 0.51844889 | 5.1225303 | pHu+ |
| 3:a | 3 | 17256775 | 0.34720312 | 4.9183282 | pHu+ |
| 3:a | 3 | 18221565 | 0.46198157 | 4.3883626 | pHu+ |
| 3:a | 3 | 18348456 | 0.07279455 | 5.2099823 | pHu+ |
| 3:a | 3 | 18598504 | 0.15273550 | 5.5729773 | pHu+ |
| 3:b | 3 | 104194096 | 0.66134818 | 5.3095180 | pHu+ |
| 4:a | 4 | 63446574 | 5.64460484 | 0.9894550 | pHu- |
| 6:a | 6 | 6356771 | 0.44110600 | 4.7090271 | pHu+ |
| 13:a | 13 | 14544320 | 1.23371366 | 5.4294493 | pHu+ |
| 14:a | 14 | 1718020 | 0.21743406 | 4.7147736 | pHu+ |
| 17:a | 17 | 8225007 | 0.16401880 | 4.5066901 | pHu+ |
| 21:a | 21 | 4259090 | 0.32265976 | 4.5090274 | pHu+ |
| 22:a | 22 | 4635167 | 0.83432065 | 5.3513924 | pHu+ |

## Approximating moments of Wright-Fisher process with a Taylor expansion

### 1 Framework

Let *X*_*n*_ be a sequence of random variables following a Wright-Fisher bi-allelic allele frequency discrete process. This means that the conditional law of *X*_*n*+1_ given *X*_*n*_ is 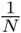 multiplied by a binomial with parameters *N* and *f* (*X*_*n*_) where *f* is a function, let’s say *f* ∈ *𝒞*^∞^([0, 1], [0, 1]) for simplicity. This is commonly written as

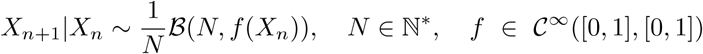

The aim of this development is to accurately approximate moments of this process without having to compute the whole process transition matrix. One can derive the following identities :

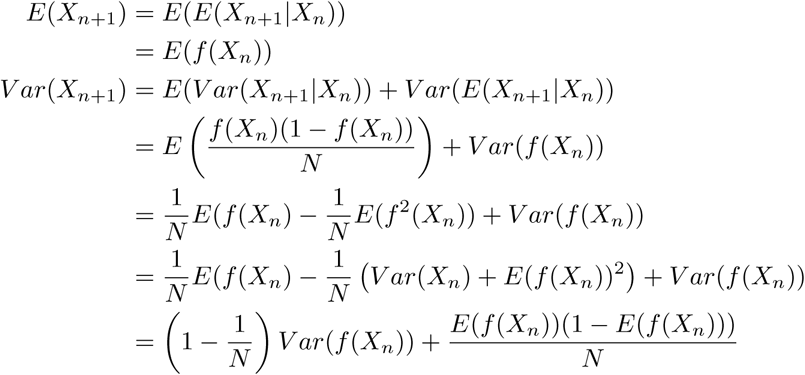

As *f* is non linear function, we can’t say that either *E*(*f* (*X*_*n*_)) = *f* (*E*(*X*_*n*_)) or *V ar*(*f* (*X*_*n*_)) = *f* ′(*E*(*X*_*n*_))*V ar*(*X*_*n*_) (which is the case when *f* is linear). However we can make an approximation around some point using a Taylor expansion. In the following, *µ*_*n*_ designate a deterministic sequence expected to be our approximation of *E*(*X*_*n*_). In the same way, let 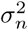 designate another deterministic sequence expected to be our approximation of *V ar*(*X*_*n*_). One can set the error terms for moments : *ε*_*n*_ = *E*(*X*_*n*_) *-µ*_*n*_ and 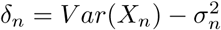

One can get the relation :

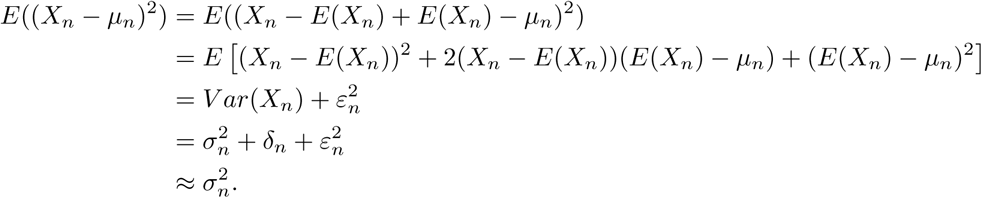

### 2 Lacerda and Seoighe (2014) approximation

Let’s do a 1^st^ taylor expansion of any function *g* around the mean of a random variable *X* :

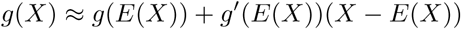

So by using mean and variance properties, one get :

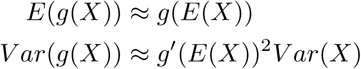

With these approximations, one get imediately the following identities :

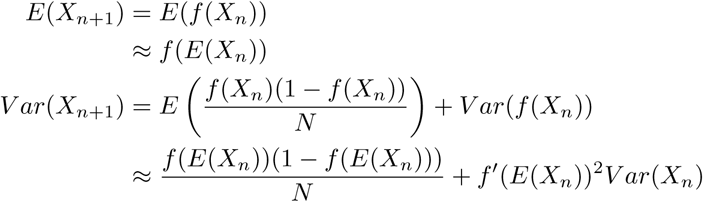

### 3 Terhorst *et al.* (2015) approximation

Let’s write *X*_*n*_ = *x*_*n*_ + *δX*_*n*_ where *x*_*n*+1_ = *f* (*x*_*n*_) and do the 2^nd^ order Taylor expansion about this quantity:

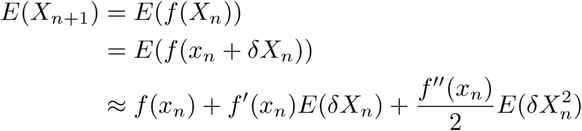

By definition, one get also :

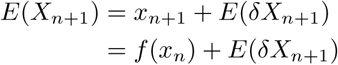

So one can obtain the *δX*_*n*_ recursion:

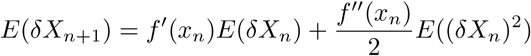

At this point, the author said :

Inductively, assuming that we can compute *E*(*δX*_*n*_) and *E*((*δX*_*n*_)^2^), this enable us to compute *E*(*X*_*n*_) and *V ar*(*X*_*n*_) = *V ar*(*δX*_*n*_). This approach was previously employed by Barton *et al.* (2005) to obtain order 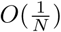 approximations to these moments. Here, we have used the same idea but automated the symbolic algebra and code generation needed to generate the recursions to higher orders of accuracy.

This suggest that the author did implement an higher order recursion. However, they have not given recursions for *E*((*δX*_*n*_)^2^), so assuming they did in the same way than for *E*(*δX*_*n*_), one obtain :

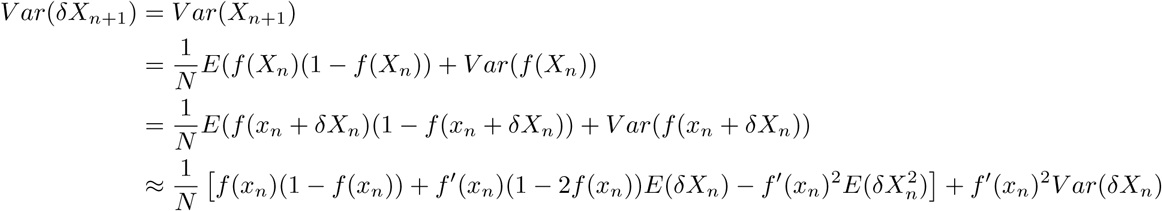

Using the fact that *V ar*(*δX*_*n*_) = *E*((*δX*_*n*_)^2^) *-E*(*δX*_*n*_)^2^, all relations needed are established

### 4 Taylor expansion

Remember the following recursions :

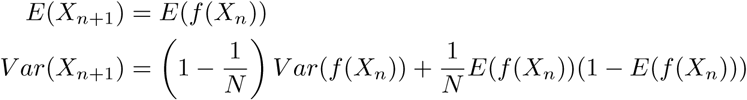

As *f* isn’t linear, *E*(*f* (*X*_*n*_)) has no closed form (the same problem occurs for higher moments). So one needs approximation for these quantities.

To do this, one can expand the *f* function around our mean approximation *µ*_*n*_ assumed to be close from *E*(*X*_*n*_). The following formula could help in that way :

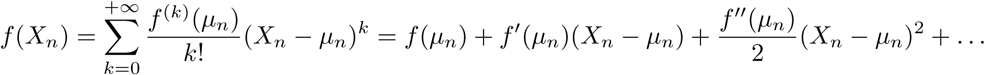

Assuming that all the operations done are legal, one can optain the following relations :

### Proposition 1

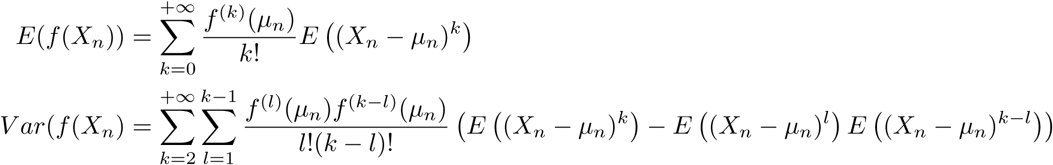

*Proof.* By taking the expectancy of 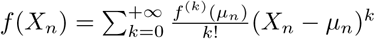 One gets immediately the 1^st^ point :

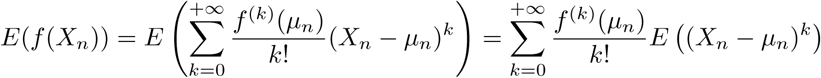

In the same way, the other relations can be obtained :

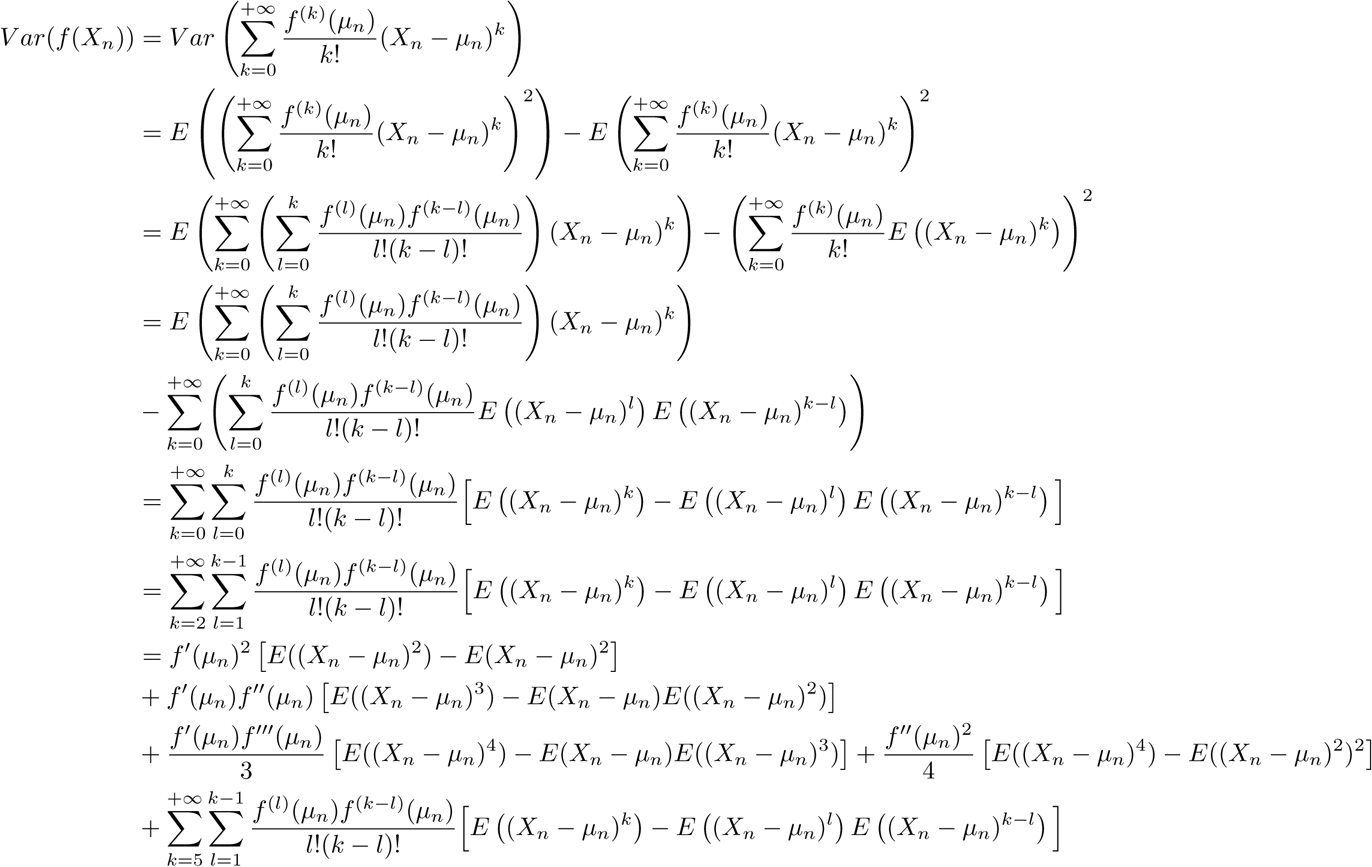

□

#### 4.1 1^st^ order approximation

In this part, the previous relations are used to pinpoint recursions between the moments approximations :

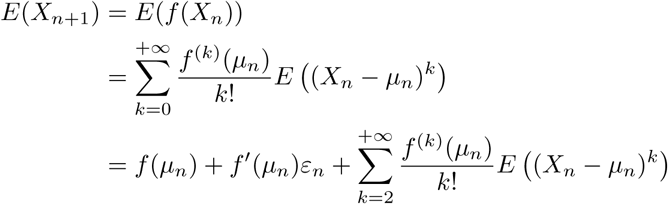

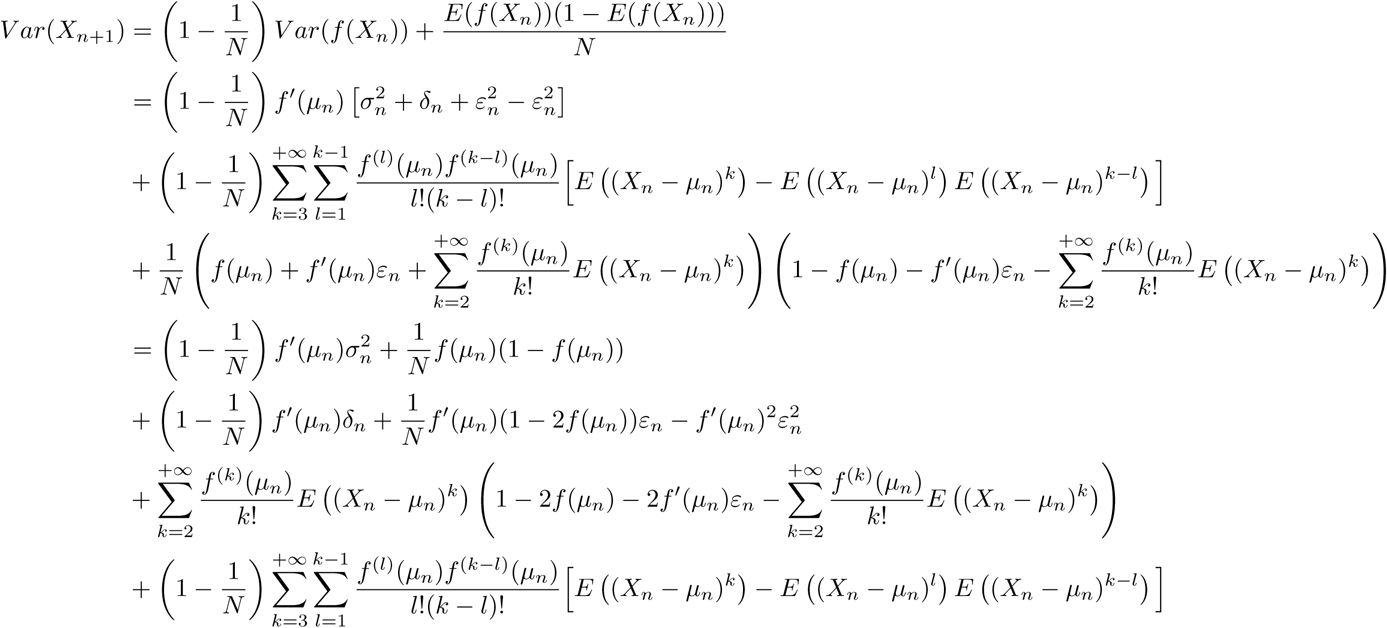

#### 4.2 2^nd^ order approximation

In this part, the previous relations are used to pinpoint recursions between the moments approximations :

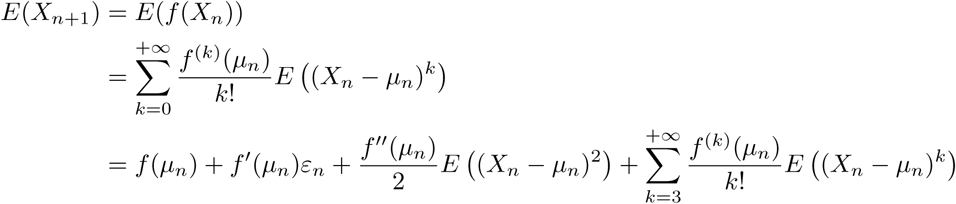

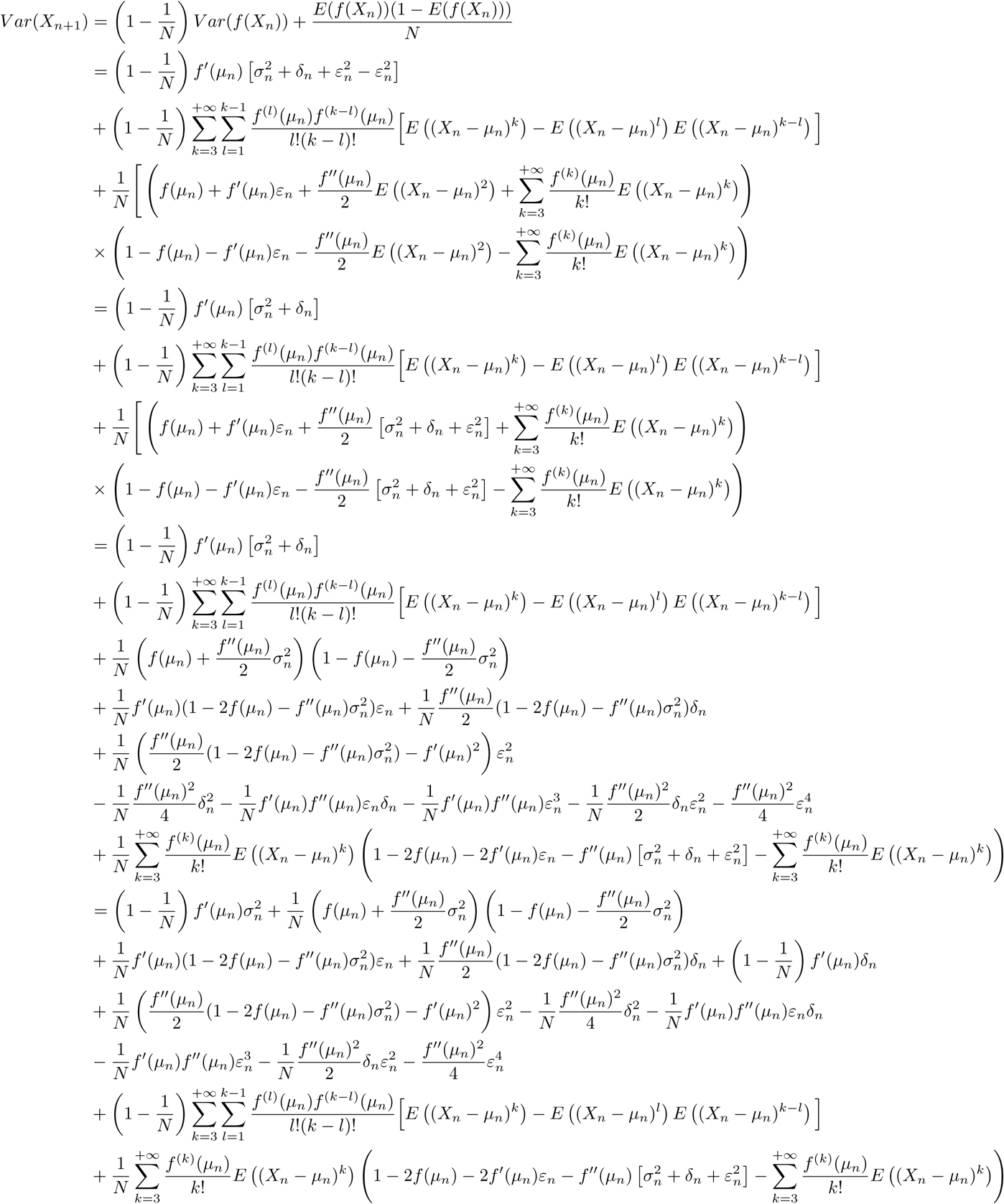

### 5 Recursions

With the relations established before, one can set different recursions schemes to approximate moments of the true Wright-Fisher process.

#### 5.1 From Lacerda and Seoighe’s derivations

One obtained the following :

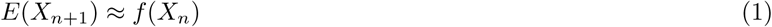

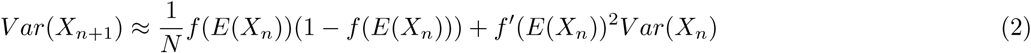

Recursions used by Lacerda and Seoighe are:

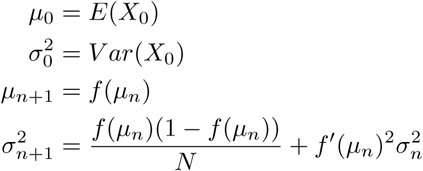

**Remark 1.** *Note that in their case, they took f having the form* :

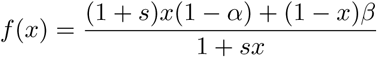

#### 5.2 From Terhorst *et al.*’s (2015) derivations

These recursions are more tricky than previous ones because it’s a five crossed sequence recursion :

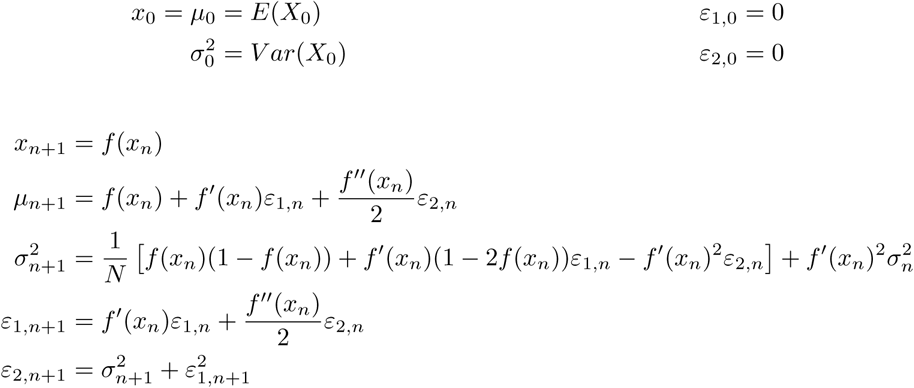

#### 5.3 From 1^st^ order approximation

In this part, let 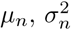 be defined from the following relations :

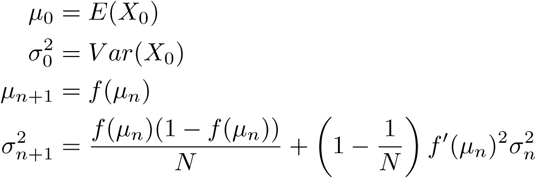

#### 5.4 From 2^nd^ order approximation

In this part, let 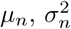 be defined from the following relations :

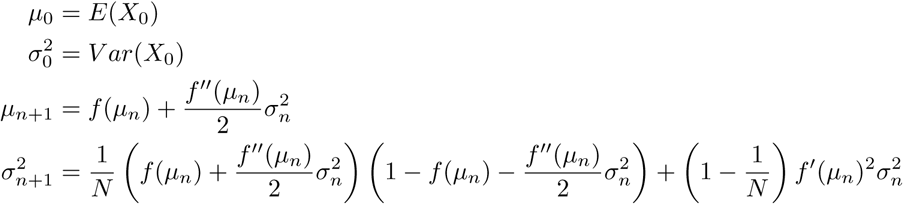

